# The WZA: A window-based method for characterizing genotype-environment association

**DOI:** 10.1101/2021.06.25.449972

**Authors:** Tom R. Booker, Sam Yeaman, Michael C. Whitlock

**Affiliations:** Department of Biological Sciences, University of Calgary, Calgary, Canada; Department of Zoology, University of British Columbia, Vancouver, Canada; Biodiversity Research Centre, University of British Columbia, Vancouver, Canada

**Keywords:** Local adaptation, population genetics, landscape genomics, GEA

## Abstract

Genotype environment association (GEA) studies have the potential to elucidate the genetic basis of local adaptation in natural populations. Specifically, GEA approaches look for a correlation between allele frequencies and putatively selective features of the environment. Genetic markers with extreme evidence of correlation with the environment are presumed to be tagging the location of alleles that contribute to local adaptation. In this study, we propose a new method for GEA studies called the weighted-Z analysis (WZA) that combines information from closely linked sites into analysis windows in a way that was inspired by methods for calculating *F_ST_*. We analyze simulations modelling local adaptation to heterogeneous environments either using a GEA method that controls for population structure or an uncorrected approach. In the majority of cases we tested, the WZA either outperformed single-SNP based approaches or performed similarly. The WZA outperformed individual SNP approaches when the measured environment is not perfectly correlated with the true selection pressure or when a small number of individuals or demes was sampled. We apply the WZA to previously published data from lodgepole pine and identified candidate loci that were not found in the original study.

## Introduction

Studying local adaptation can provide a window into the process of evolution, yielding insights about the nature of evolvability, constraints to diversification, and the how the interplay between a species and its environment shapes its genome (e.g. Savolainen 2013). Understanding local adaptation can also benefit practical applications such as in forestry where many species of economic interest exhibit pronounced trade-offs in fitness across environments. Characterizing such trade-offs may help identify alleles involved in local adaptation, revealing candidate genes important for breeding or informing conservation management programs for buffering against the consequences of anthropogenic climate change (Aitken and Whitlock 2013). Whatever the aim or application, a first step in studying the basis of local adaptation is to identify the genes that are driving it.

A potentially powerful method for identifying the genomic regions involved in local adaptation is genotype-environment association (GEA) analysis, which has been widely adopted in recent years. Alleles may vary in frequency across a species’ range in response to local environmental conditions that give rise to spatially varying selection pressures (Haldane 1948). For that reason, genetic variants that exhibit strong correlations with putatively selective features of the environment are often interpreted as a signature of local adaptation (Coop et al. 2010). Genotype-environment association (GEA) studies examine such correlations. Allele frequencies for many genetic markers, typically single nucleotide polymorphisms (hereafter SNPs), are estimated in numerous locations across a species’ range. Correlations between allele frequency and environmental variables are calculated then contrasted for sites across the genome. It is assumed in GEA studies that current heterogeneity in the environment (whether biotic or abiotic) reflects the history of selection.

Numerous approaches for performing GEA analyses have been proposed. If individuals are sequenced, GEA can be performed by regressing environments on genotypes as a form of genome-wide association study, for example using the *GEMMA* package (Zhou et al. 2013). However, to estimate SNP effects with reasonable statistical power, many individuals may need to be sequenced. A cost-effective alternative is pooled sequencing (hereafter pooled-seq), where allele frequencies for populations of individuals are estimated rather than individual genotypes (Schlötterer et al. 2014). In this study, we focus on analyses that can be performed on pooled-seq datasets given the wide adoption of that protocol in the GEA literature.

The most straightforward way to perform a GEA analysis is to simply examine the correlation between allele frequencies and environmental variables measured in multiple populations, for example using rank correlations such as Spearman’s *ρ* or Kendall’s *τ*. This simple approach may commonly lead to false positives, however, if there is environmental variation across the focal species’ range that is correlated with patterns of gene flow or historical selection (Meirmans 2012; Novembre and Di Rienzo 2009). For example, consider a hypothetical species inhabiting a large latitudinal range. If this species had restricted migration and exhibited isolation-by-distance, neutral alleles may be correlated with any environmental variable that happened to correlate with latitude, as population structure would also correlate with latitude.

Several approaches have been proposed to identify genotype-environment correlations above and beyond what is expected given an underlying pattern of population structure and environmental variation. For example, the commonly used *BayPass* package (Gautier 2015), an extension of *BayEnv* by Coop et al. (2010), estimates correlations between alleles and environmental variables in a two-step process. First, a population covariance matrix (**Ω**) is estimated from SNP data. Second, correlations between the frequencies of individual SNPs and environmental variables are estimated treating **Ω** in a manner similar to a random effect in a generalized mixed model. In a recent study, Lotterhos (2019) compared several of the most commonly used packages for performing GEA on pooled-seq datasets; including *BayPass* (Gautier 2015), latent-factor mixed models (LFMMs) as implemented in the LEA package (Frichot et al. 2013; Frichot and François 2015), redundancy analysis (RDA; see Forester et al. 2016, 2018) and a comparatively simple analysis calculating Spearman’s *ρ* between allele frequency and environment. Of the methods they tested, Lotterhos (2019) found that the GEA approaches that did not correct for population structure (i.e., Spearman’s *ρ* and RDA) had higher power to detect local adaptation compared to *BayPass* or LFMMs. In their standard application to genome-wide datasets, all the GEA analysis methods mentioned provide a summary statistic for each marker or SNP.

Individual SNPs may provide very noisy estimates of summary statistics, but closely linked SNPs are not independently inherited and may have highly correlated evolutionary histories. As a way to reduce noise, genome scan studies often aggregate data across adjacent markers into analysis windows based on a fixed physical or genetic distance or number of SNPs (Hoban et al. 2016). In the case of *F_ST_*, the standard measure of population differentiation, there are numerous methods for combining estimates across sites (see Bhatia et al. (2013)). In Weir and Cockerham’s (1984) method, for example, estimates of *F_ST_* for individual loci are combined into a single value with each marker’s contribution weighted by its expected heterozygosity.

In the context of GEA studies, each marker or SNP provides a test of whether a particular genealogy is correlated with the pattern of environmental variation. In the extreme case of a non-recombining region, all SNPs would share the same genealogy and thus provide multiple tests of the same hypothesis. For recombining portions of the genome, however, linked sites will not have the same genealogy, but genealogies may be highly correlated. Similar to combining estimates of *F_ST_* to decrease statistical noise, combining GEA tests performed on individual markers may increase the power of GEA studies to identify genomic regions that contribute to local adaptation.

In this study, we propose a general method for combining the results of single SNP GEA scores into analysis windows that we call the weighted-Z analysis (WZA), and test its efficacy using simulations. We generate datasets modelling a pooled-sequencing experiment where estimates of allele frequency are obtained for numerous populations across a species’ range. Using our simulated data, we compare the performance of the WZA to Kendall’s *τ* (because Lotterhos (2019) found that this method had high power) as well as *BayPass* (Gautier 2015), as it is a widely used approach that corrects for population structure in GEA studies. Additionally, we compare the WZA to another window-based GEA approach proposed by Yeaman et al. (2016). We found that the WZA is particularly useful when GEA analysis is performed on small samples and when results for individual SNPs are statistically noisy. We re-analyze previously published lodgepole pine (*Pinus contorta*) data using the WZA and find several candidate loci that were not identified using the methods of the original study.

## The Weighted-*Z* Analysis

In this study, we propose the Weighted-*Z* Analysis (hereafter, the WZA) for combining information across linked sites in the context of GEA studies. Specifically, we aim to combine information from multiple SNPS within the same small genomic region to ask whether that region shows associations between its local allele frequencies and local environment.

The WZA uses the weighted-*Z* test from the meta-analysis literature that combines *p*-values from multiple independent hypothesis tests into a single score (Mosteller and Bush 1954; Liptak 1958; Stouffer et al. 1949). In the weighted-*Z* test, each of the *n* independent tests is given a weight that is proportional to the inverse of its error variance (Whitlock 2005). We use the expected heterozygosity of each SNP in a gene or window for the weights in the WZA, following Weir and Cockerham (1984), as their classic method performs well in a similar evolutionary context, where the aim is to quantify divergence in allele frequencies among populations. At a given polymorphic site, we denote the average frequency of the minor allele across populations as *p̅* (*q̅* corresponds to the frequency of the major allele). Sites with higher values of *p̅* q̅ (will carry more information about the underlying genealogy.

We combine information about genetic correlations with the environment from biallelic markers (typically SNPs) present in a focal genomic region into a single weighted-*Z* score (*Z_w_*). The genomic region in question could be a gene or genomic analysis window. For each SNP with a minor allele frequency greater than 0.05 in the genomic window, we measure the association between the SNP’s local allele frequency and the local environment in some way and use the *p*-value of a test of no association for each SNP. (The exact measure used here may vary; in this paper we test the use of two such measures, described below.)

These *p-*values from each SNP in a window are combined using Stouffer’s weighted *Z* approach. We calculate *Z_W,K_* for genomic region *k*, which contains *n* SNPs, as

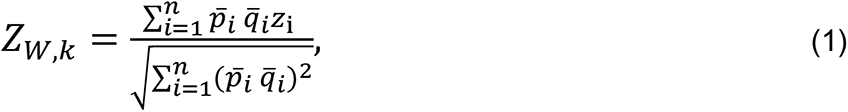

where *p̅_i_* is the mean allele frequency across populations and *z_i_* is the standard normal deviate calculated from the one-sided *p*-value for SNP *i*. A given *p*-value can be converted into a *z_i_* score by finding the corresponding quantile of the standard normal distribution, for example using the *qnorm* function in R.

When we apply the WZA in this study, we compared two different statistics as input: empirical *p*-values calculated from the genome-wide distribution of parametric *p*-values from Kendall’s *τ* correlating the local environmental variable and local allele frequency (referred to as WZA*τ*), and empirical *p*-values calculated from the genome-wide distribution of Bayes factors as obtained using the *BayPass* program (referred to as WZA_BP_; see below).

Under the null hypothesis that there is no correlation between allele frequency and environment and no spatial population structure, the expected distribution of correlation coefficients in a GEA would be normal about 0, with a uniform distribution of *p*-values. However, as will often be the case in nature, there may be an underlying correlation between population structure and environmental variation that will cause these genome-wide distributions to deviate from this null expectation. The average effect of population structure on individual SNP scores can be incorporated into an analysis by converting an individual SNP’s squared correlation coefficient or parametric *p*-value into empirical *p*-values based on the genome-wide distribution (following the approach of Hancock et al. [2011]). To calculate empirical *p*-values, we rank all values (from smallest to largest in the case of *p-*values) and divide the ranks by the total number of tests performed (i.e. the number of SNPs or markers in the analysis window). Note that in practice, we calculated empirical *p*-values after removing SNPs with minor allele frequency less than 0.05 and would recommend that others perform similar filtering. In empirical studies with varying levels of missing data across the genome, it may be preferable to rank the parametric *p*-values rather than the correlation coefficients themselves as there may be varying power to calculate correlations across the genome. With the empirical *p*-value procedure, aggregating information using the WZA will identify genomic regions with a pattern of GEA statistics that deviate from the average genome-wide. A feature of the WZA is that many tests can potentially be used as input as long as individual *p*-values provide a measure for the strength of evidence against a null hypothesis.

## Materials and Methods

In the previous section we described the mechanics of our new method, the WZA. The rest of this paper is devoted to a test of the relative efficacy of the WZA compared to other widely used approaches. Note that Lotterhos (2019) identified a simple rank correlation on individual SNPs as having among the highest power of the GEA analyses that have been tested, making such a method a good standard of comparison, and the most common GEA method used is BayPass (Gautier 2015). We use these two existing methods as our baseline of comparison for WZA.

To do these tests, we will simulate populations evolving on a variety of different environmental landscapes, with the selective optima varying over space. We use relatively weak selection, so that we are simulating the most difficult loci to find with a GEA. The present section describes the simulation conditions we used for these tests.

### Simulating local adaptation

We performed forward-in-time population genetic simulations of local adaptation to determine how well the WZA was able to identify the genetic basis of local adaptation. GEA studies are often performed on large spatially extended populations that may be comprised of hundreds of thousands of individuals. However, it is computationally infeasible to model selection and linkage in long chromosomal segments (>1Mbp) for such large populations. For that reason, we simulated relatively small populations containing 19,600 diploid individuals in total and scaled population genetic parameters to model a large population. We based our choice of population genetic parameters on estimates for conifer species. A representative set of parameters is given in Table S1 and in the Appendix we give a breakdown and justification of the parameters we chose. All simulations were performed in *SLiM* v3.4 (Messer and Haller 2019).

We simulated meta-populations inhabiting and adapting to heterogeneous environments and modelled the population structure on an idealized conifer species. In conifers, strong isolation-by-distance has been reported and overall mean *F_ST_* < 0.10 has been estimated in several species (Mimura and Aitken 2007; Mosca et al. 2014). We thus simulated individuals inhabiting a 2-dimensional stepping-stone population made up of 196 demes (i.e. a 14 × 14 grid). Each deme consisted of *N_d_* = 100 diploid individuals. We assumed a Wright-Fisher model so demes did not fluctuate in size over time. Migration was limited to neighboring demes in the cardinal directions and the reciprocal migration rate between demes (*m*) was set to 0.0375 in each possible direction to achieve an overall *F_ST_* for the metapopulation of around 0.04 (Figure S1). As expected under restricted migration, our simulations exhibited a strong pattern of isolation-by-distance (Figure S1). Additionally, we simulated metapopulations with no spatial structure (i.e., finite island models). In these simulations, we used the formula

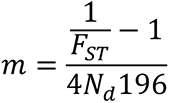

(Charlesworth and Charlesworth 2010; pp319) to determine that a migration rate between each pair of demes of *m* = 4.12 x 10^-4^ would give a target *F_ST_* of 0.03.

The simulated organism had a genome containing 1,000 genes uniformly distributed onto 5 chromosomes. We simulated a chromosome structure in *SLiM* by including nucleotides that recombined at *r* = 0.5 at the hypothetical chromosome boundaries. Each chromosome contained 200 segments of 10,000bp each. We refer to these segments as genes for brevity, although we did not model an explicit exon/intron or codon structure. It has been reported that linkage disequilibrium (LD) decays rapidly in conifers, with LD between pairs of SNPs decaying to background levels within 1,000bp or so in several species (Pavy et al. 2012). In our simulations, recombination within genes was uniform and occurred at a rate of *r* = 10^-7^ per base-pair, giving a population-scaled recombination rate (4*N_d_r*) of 0.0004. The recombination rate between the genes was set to 0.005, effectively modelling a stretch of 50,000bp of intergenic sequence. Given these recombination rates, LD decayed rapidly in our simulations with SNPs that were approximately 600bp apart having, on average, half the LD of immediately adjacent SNPs in neutral simulations (Figure S1). Thus, patterns of LD decay in our simulations were broadly similar to the patterns reported for conifers.

We incorporated spatial variation in the environment into our simulations using a discretized map of degree days below 0 (DD0) across British Columbia (BC). We generated the discretized DD0 map by first downloading the map of DD0 for BC from ClimateBC (http://climatebc.ca/; Wang et al. 2016; Figure 1A). Using Dog Mountain, BC as the reference point in the South-West corner (Latitude = 49.37, Longitude = −122.97), we extracted data in a rectangular grid with edges 3.6 degrees long in terms of both latitude and longitude, an area of approximately 266 × 400*km*^2^ (Figure 1A). We divided this map into a 14 × 14 grid, calculated the mean DD0 scores in each grid cell, converted them into standard normal deviates (i.e. *Z*-scores) and rounded up to the nearest third. We used the number of thirds of a *Z*-score as phenotypic optima in our simulations. We refer to this map of phenotypic optima as the *BC* map (Figure 1B).

**Figure 1.**
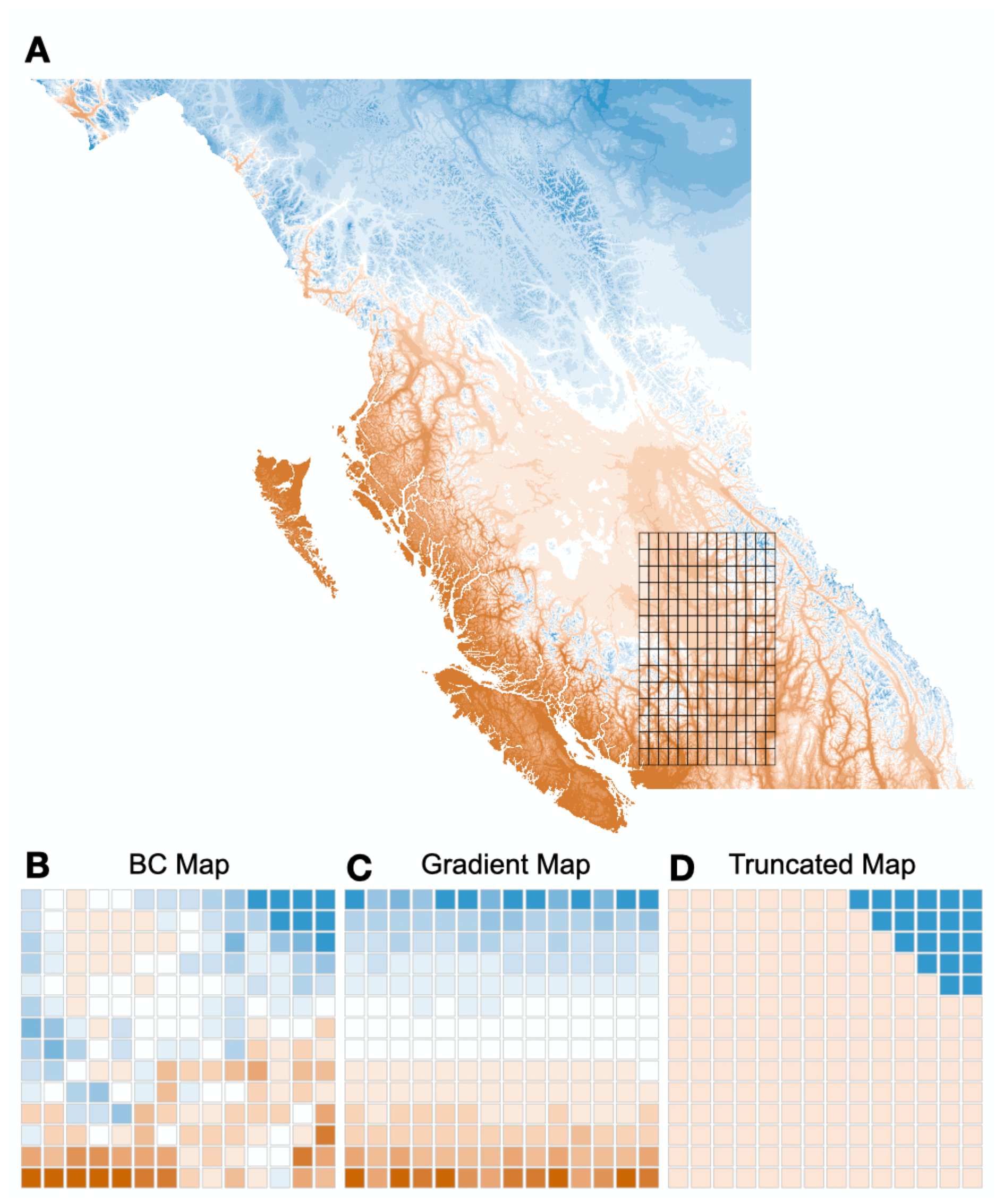
A) Degree days below zero across British Columbia, the overlain grid in A shows the locations we used to construct phenotypes for our simulated populations. B) A discretized map of DD0 in Southern British Columbia, we refer to the map in B as the *BC map*. C) A 1-dimensional gradient of phenotypic optima, we refer to this as the Gradient map. D) A model of selection acting on a small proportion of the population, we refer to this map as the Truncated map.

We used data from the *BC* map to generate two additional maps of environmental variation. First, we ordered the data from the *BC* map along one axis of the 14 × 14 grid and randomised optima along the non-ordered axis. We refer to this re-ordered map as the *Gradient* map (Figure 1C). Second, we generated a map where selection differed over only a small portion of the environmental range. For some species, fitness optima may differ only beyond certain environmental thresholds (*e.g.* temperature above vs. below 0°C), leading to a non-normal distribution of phenotypic optima. To model such a situation, we set the phenotypic optimum of 20 demes in the top-right corner of the meta-population to +3 and set the optimum for all other populations to –1. We chose 20 demes as it represented approximately 10% of the total population. We refer to this map as the *Truncated* map (Figure 1D).

We simulated local adaptation using models of either directional or stabilizing selection. In both cases, there were 12 causal genes distributed evenly across four simulated chromosomes that potentially contributed to local adaptation. With directional selection, mutations affecting fitness could only occur at a single nucleotide position in the center of the 12 potentially selected genes. Directionally selected mutations had a spatially antagonistic effect on fitness. In deme *d* with phenotypic optimum *θ_d_*, the fitness of an individual homozygous for the selected allele was 1 + *s_a_θ_d_*(selected alleles were semi-dominant). The fitness affecting alleles had a mutation rate of 3 × 10^-7^ in simulations modelling directional selection and a fixed *s_a_* = 0.003 (see *Appendix*).

Under stabilizing selection, the mutations that occurred in the 12 genes had a normal distribution of phenotypic effects, with variance 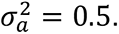 Phenotype-affecting mutations occurred at a rate of 10^-7^ per base-pair in the 12 genes, and could occur at any of the 10,000 sites within a given gene. An individual’s phenotype was calculated as the sum of the effects of all phenotype-affecting mutations. We calculated an individual’s fitness using the standard expression for Gaussian stabilizing selection,

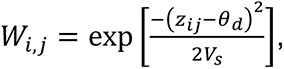

where *z_i,j_*. is the phenotype of the *i*^th^ individual in environment *j* and *V_s_* is the variance of the Gaussian fitness function (Walsh and Lynch 2018). We set *V_s_*= 196 so that there was a 40% fitness difference between individuals perfectly adapted to the two extremes of the distribution of phenotypic optima. This was motivated by empirical studies of local adaptation that have demonstrated such fitness differences in numerous species (Hereford 2009; Bontrager et al. 2020); see *Appendix*.

We ran simulations for a total of 200,102 generations. The 19,600 individuals initially inhabited a panmictic population that evolved neutrally. After 100 generations, the panmictic population divided into a 14 × 14 stepping-stone population and evolved either strictly neutrally (when modelling directional selection) or with a phenotypic optimum of 0 for all demes (when modelling stabilizing selection). After 180,000 generations, we imposed the various maps of phenotypic optima and simulated for a further 20,000 generations. For selected mutations, we used the “*f*” option for *SLiM*’s mutation stack policy, so only the first mutational change was retained. Using the tree-sequence option in *SLiM* (Haller et al. 2019), we tracked the coalescent history of each individual in the population. At the end of each simulation, neutral mutations were added at a rate of 10^-8^ using *PySLiM* (https://pyslim.readthedocs.io/en/latest/). For each combination of map and mode of selection, we performed 20 replicate simulations.

### Classifying simulated genes as locally adapted

To evaluate the performance of different GEA methods, we needed to identify which of the 12 causal genes contributed to local adaptation and which did not in each replicate of our simulated data. As described above, our simulations incorporated a stochastic mutation model so from replicate to replicate the genes that contributed to local adaptation varied and, in the case of stabilizing selection, so did the effect size of the alleles in those genes.

For simulations modelling directional selection, we identified locally adapted genes based on the mean fitness of their alleles at the single variable site in each gene with a polymorphism. Our measure of local adaptation was the covariance between the mean fitness contributed by the selected allele in each population and the environment.

For simulations modelling stabilizing selection, we identified locally adapted genes based on the covariance of the environment and the phenotypic effects of their alleles, summed across all variant sites within each gene. For a given gene, we summed the additive phenotypic effects of all non-neutral variants and took the average for each population. Our measure of local adaptation for each gene was the covariance between that average additive phenotypic effect and environmental variation (we refer to this as *Cov(Phen, Env*)).

For both selection regimes, we defined locally adapted genes as those with a covariance between environment and allelic effect (in fitness or phenotypic terms) greater than 0.005. When modelling directional selection, an average of 6.35, 6.50 and genes (out of 12) contained genetic variants that established and contributed to local adaptation for the *BC map*, the *Gradient map* and the *Truncated map*, respectively. In our simulations modelling stabilizing selection, individuals’ and population mean phenotypes closely matched the phenotypic optima of their local environment (Figure S2). The average numbers of genes contributing to local adaptation in individual replicates in these simulations were 7.15, 6.45 and 5.35 for the *BC map*, the *Gradient map* and the *Truncated map*, respectively. However, when analyzing stabilizing selection simulations, we calculated the proportion of the total *Cov(Phen, env)* explained by a particular set of genes rather the number of true positives.

### Analysis of simulation data

We performed GEA on our simulated data using either Kendall’s *τ*-b (hereafter Kendall’s *τ*), a rank correlation that does not model population structure, or *BayPass*, which corrects for a population covariance matrix (Gautier 2015). For all analyses, except where specified, we analyzed data for a set of 40 randomly selected demes and sampled 50 individuals from each to estimate allele frequencies. We sampled individuals from the same set of demes for all analyses, shown in Figure S3. Each simulation replicate included 1,000 genes, and after excluding alleles with a minor allele frequency less than 0.05 there was an average of 23.3 SNPs per gene. We ran *BayPass* following the “worked example” in section 5.1.2 of the manual provided with the software.

We used three different methods to summarize the GEA results for each gene in each simulation replicate: a single SNP-based approach, the WZA and the top-candidate method developed by Yeaman et al. (2016). For all three tests, we used either the *p-* values from Kendall’s *τ* or Bayes factors from *BayPass*.

- For the implementation of the single SNP-based approach, the SNPs with the most extreme test statistic (i.e. smallest *p*-value or largest Bayes factor) for each gene were recorded and other SNPs in the gene were subsequently ignored. This was done to prevent multiple outliers that are closely linked from being counted as separate hits. The single-SNP based method is perhaps most similar to how GEA analyses are typically interpreted, as it relies upon the evidence from the most strongly associated SNP to assess significance for a closely linked gene.
- We implemented a simplified version of the top-candidate method proposed by Yeaman et al. (2016). The top-candidate method attempts to identify regions of the genome involved in local adaptation under the assumption that such regions may contain multiple sites that exhibit strong correlation with environmental variables. The top-candidate method asks whether there is a significant excess of “outlier” SNPs in a region compared to what one would expect given the genome wide distribution. The number of outliers in each genomic region is compared to the expected number of outliers based on the genome-wide proportion of SNPs that are outliers, using a binomial test. We defined outliers as those within the 99^th^ percentile of scores genome wide. The *p*-value from the binomial test is used as a continuous index.
- For the implementation of the WZA, we converted the *p*-values (from Kendall’s *τ*) or Bayes factors (from *BayPass*), into empirical *p*-values. For each of the *n* SNPs present in a gene, empirical *p*-values were converted into *z* scores and used to calculate WZA scores using Equation 1.

We examined the effect of variation in recombination on the properties of the WZA by manipulating the tree-sequences that we recorded in *SLiM*. In our simulations, genes were 10,000 bp long, so to model genomic regions of low recombination rate, we extracted the coalescent trees that corresponded to the central 1,000bp or 100bp of each gene. For the 1,000bp and 100bp intervals, we added mutations at 10× and 100× the standard mutation rate, respectively.

All SNPs present in each 10,000bp gene in our simulations were analyzed together. However, to explore the effect of window size on the performance of the WZA, we calculated WZA scores for variable numbers of SNPs. In these cases, we calculated WZA scores for all adjacent sets of *g* SNPs and retained the maximum WZA score for all sets of SNPs in the gene.

Tree sequences were manipulated using the *tskit* package. Mutations were added to trees using the *msprime* (Kelleher et al. 2016;

https://tskit.dev/msprime/docs/stable/intro.html), *tskit* and *PySLiM* workflow (https://pyslim.readthedocs.io/en/latest/). *F_ST_* and *r*^2^ (an estimator of linkage disequilibrium) were calculated using custom Python scripts that invoked the *scikit-allel* package (https://scikit-allel.readthedocs.io/en/stable/).

### Analysis of data from lodgepole pine

We re-analyzed a previously published population genomic dataset for lodgepole pine, *Pinus contorta*, a conifer that is widely distributed across the Northwest of North America. Briefly, Yeaman et al. (2016) collected samples from 254 populations across British Columbia and Alberta, Canada and Northern Washington, USA. The lodgepole pine genome is very large (approximately 20Gbp), so Yeaman et al. (2016) used a sequence capture technique based on the *P. contorta* transcriptome. Allele frequencies were estimated for many markers across the captured portion of the genome by sequencing 1-4 individuals per population. Yeaman et al. (2016) performed GEA on each SNP using Spearman’s *ρ* and used their top-candidate method (see above) to aggregate data across sites within genes. We downloaded the data for individual SNPs from the Dryad repository associated with Yeaman et al. (2016) (https://doi.org/10.5061/dryad.0t407). We converted Spearman’s *ρ p*-values into empirical *p*-values and performed WZA on the same genes analyzed by Yeaman et al. (2016). We also repeated the top-candidate method, classifying SNPs with empirical *p-values* < 0.01 as outliers. However, as above, we use the *p*-value from the top-candidate method as a continuous index.

### Data and Code Availability

The simulation configuration files and code to perform the analysis of simulated data and generate the associated plots are available at *github/TBooker/GEA/WZA*. Analyses were performed using a combination of R and Python. All plots were made using *ggplot2* (Wickham 2016). Tree-sequence files for the simulated populations will be made available at Dryad and all processed GEA files are available on (*details to be determined post-submission*).

## Results

### The statistical properties of the WZA

To assess the statistical properties of the WZA, we first performed GEA analyses on populations that were evolving neutrally. Figure 2A shows the distribution of WZA*τ* scores for stepping-stone populations simulated under neutrality. The null expectation for WZA scores is the standard normal distribution (mean of 0 and standard deviation of 1), but we found that the distribution of WZA*τ* scores deviated slightly from this even under neutrality, where the mean and standard deviation of WZA*τ* scores from individual simulation replicates were approximately 0.089 and 1.38, respectively.

**Figure 2.**
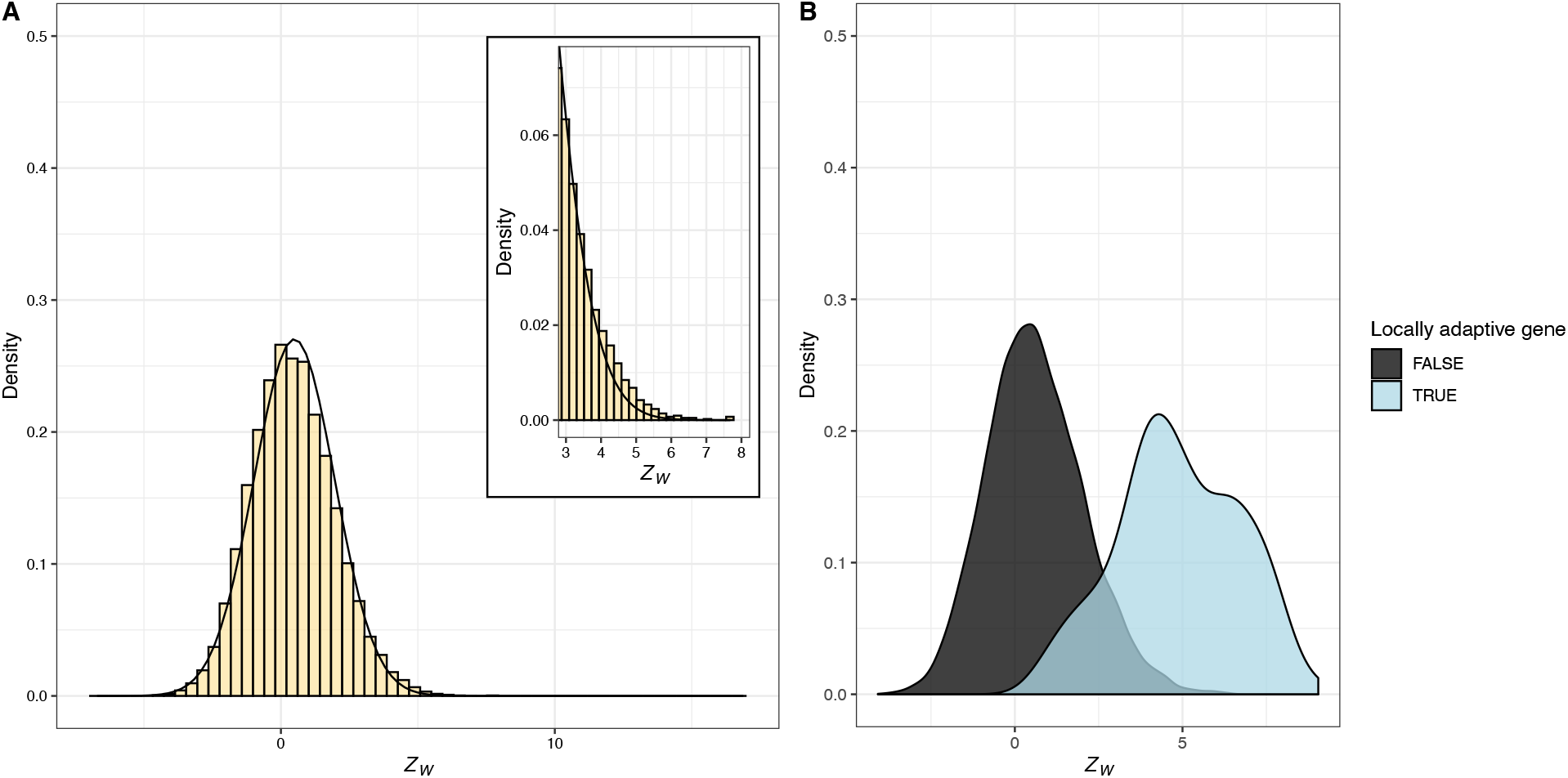
The distribution of WZA scores under neutrality and a model of local adaptation. A) A histogram of WZA*τ* scores under strict neutrality across a set of 20 replicate simulations, inset is a close-up view of the upper tail of the distribution of Z_W_ scores. The black line indicates the standard normal distribution. B) A density plot showing the separation of WZA*τ* scores for genes that are locally adaptive versus evolving neutrally across the genome of 20 simulation replicates. GEA was performed on 40 demes sampled from the *BC Map*.

Additionally, the inset histogram in Figure 2A shows that distribution of WZA*τ* scores had a somewhat thicker right-hand tail than expected under the normal distribution. A similar deviation from normality was observed when data were simulated under an island model, or when WZA was calculated using Bayes factors (Figure S4).

The deviation from the standard normal distribution is driven by non-independence of SNPs within the analysis windows we used to calculate WZA*τ* scores. To demonstrate this, we re-calculated WZA*τ* scores, but permuted the locations of SNPs across the genome, effectively erasing the signal of linkage within genes. The distribution of WZA*τ* scores in this permuted dataset closely matched the null expectation and did not have a thick right-hand tail (Figure S4; shuffled); each of 20 simulation replicates had a mean WZA*τ* indistinguishable from 0 with a standard deviation very close to 1. It is worth noting that we modelled populations that did not change in size over time. Non-equilibrium population dynamics such as population expansion may influence the distribution of WZA scores.

When evolution includes selection, WZA can often clearly distinguish regions of the genome containing loci that contribute to local adaptation from those that do not. Figure 2B shows separation of WZA*τ* scores for genes that contribute to local adaptation from those that are evolving neutrally (similar results were found for both the *Gradient* and *Truncated* maps; Figure S5). The distributions of WZA*τ* scores for locally adapted genes when modelling stabilizing selection was broader than when modelling directional selection (Figure S5), consistent with differences in the distributions of effect size for the genes involved in local adaptation under the two selection models (Figure S6). The separation of the distributions of WZA*τ* scores for locally adaptive genes versus neutrally evolving genes indicates that it may be a powerful method for identifying the genetic basis of local adaptation. The use of *p̅* q̅(as weights in the WZA improved performance over an unweighted version of the test (Figure S7A).

### Comparison of the WZA with other GEA approaches

We compared WZA to two other methods for identifying genomic regions that contribute to local adaptation from GEA data (Figure 3). To assess the performance of the different methods, we examined the top 1, 2, 3, … 50 genes in terms of WZA*τ* scores, −*log*_10_ (*p*-values) from the top-candidate method, or the single SNP Kendall’s *τ* approach. We calculated the proportion of all true positives that were identified in each case. In our simulations, there were 1,000 genes in total with approximately 6 locally adapted genes in each replicate (see Methods). For visualization purposes, we include Figure S8, which shows the –*log*_10_(*p*-values) from Kendall’s *τ* represented as a Manhattan plot for individual simulation replicates as well as WZA*τ* and top-candidate scores calculated from those data. Figure 3 compares the performance of the GEA methods across the three different maps of environmental variation that we simulated. For each of the three maps we simulated, we analyzed samples of 10, 20 or 40 demes where allele frequencies were estimated from 50 individuals sampled in each location.

**Figure 3.**
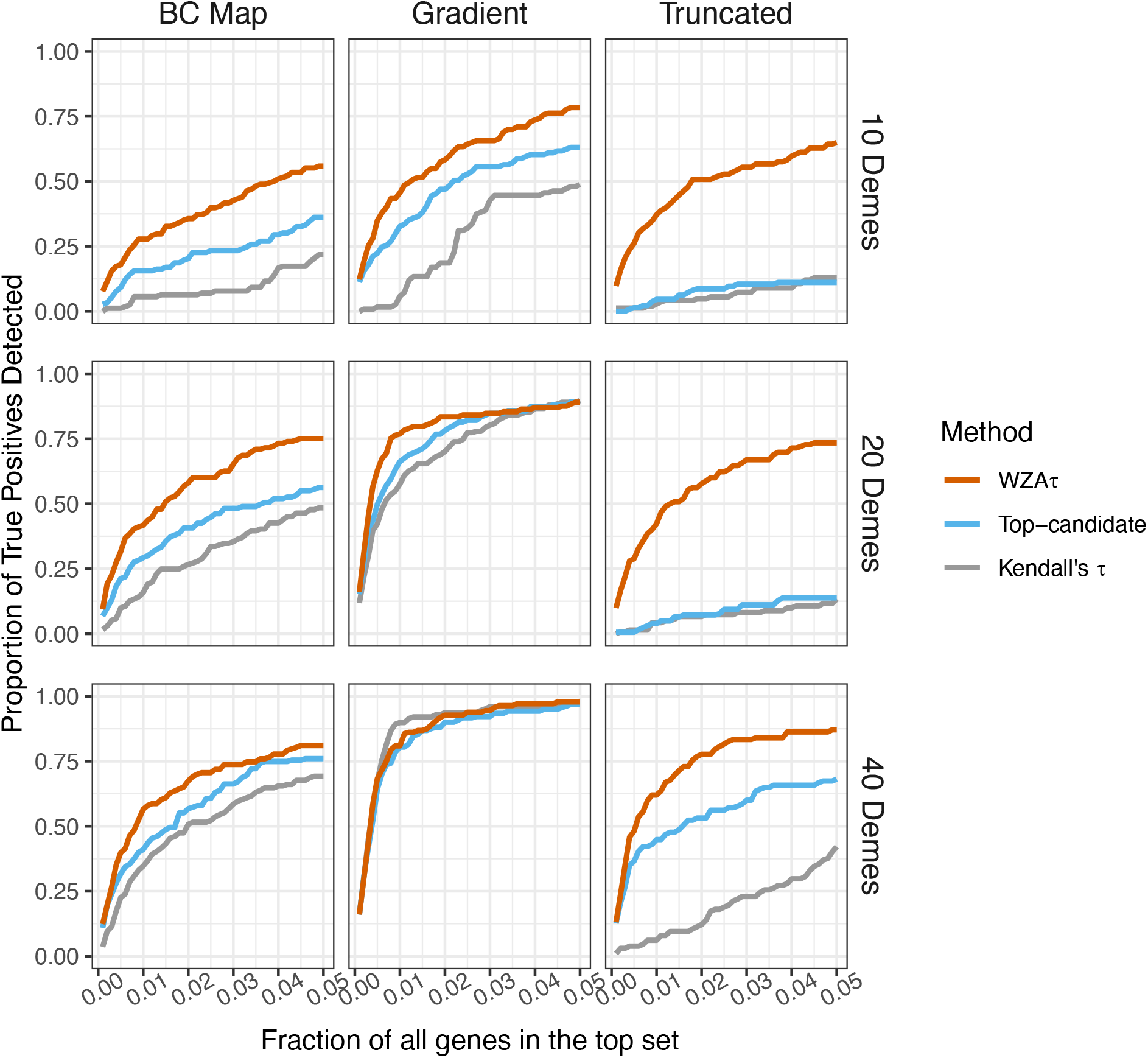
The efficacy of three GEA methods based on simulations modelling local adaptation via directional selection. In each case and separately for each method, genes were ranked in descending order of evidence for association between allele frequency and environment, and the genes with the strongest evidence for local adaptation were retained in a “top set”. The x-axes indicate the fraction of the genes which were retained in this top set. The y-axes indicate the proportion of the genes that truly contributed to local adaptation which were found in this top set. Larger values indicate a more effective method. The rows of the plot show results obtained from samples of 10, 20 or 40 demes as indicated by the labels on the right-hand side. Lines represent the means of 20 simulation replicates. In these simulations 50 individuals were sampled for each of the included populations.

Figure 3 shows that WZA*τ* substantially outperformed both the top-candidate and single SNP-based Kendall’s τ analyses in most cases. When analyzing simulations that used the *BC* map or the *Truncated* map, WZA*τ* always outperformed the top-candidate and SNP-based methods, but particularly so when fewer demes were sampled (Figure 3). When simulations assumed the *Gradient* map, WZA*τ* outperformed the other GEA methods when the sample was restricted to 10 demes, but with larger samples, the tests were more similar (Figure 3). This suggests that WZA*τ* is a powerful method for identifying regions of the genome that contribute to local adaptation in empirical analyses, but particularly so when they are performed on small samples.

An additional source of variation in GEA studies comes from the number of individuals sampled in each location. We also examined the effect that reduced sampling of individuals within each deme had on the performance of the methods. Figure S9 shows that the WZA outperforms the top-candidate and SNP-based methods when a small number of individuals is used to estimate allele frequencies.

### Effects of population structure correction

In each of the maps of environmental variation that we simulated, there was a strong correlation between environmental variables and gene flow. There was also a strong pattern of isolation-by-distance in our simulated populations (Figure S1). These two factors may make it difficult to identify genes involved in local adaptation in GEA studies (Meirmans 2012).

We compared the performance of the WZA to a widely adopted method for performing GEA that corrects for the confounding effects of population structure, *BayPass* (Gautier 2015). In all cases, WZA performed as well, or better than, *BayPass* (Figure 4). WZA performed much better than *BayPass* when selection was directional, but WZA was also significantly more likely to identify the genes underlying local adaptation with stabilizing selection.

**Figure 4.**
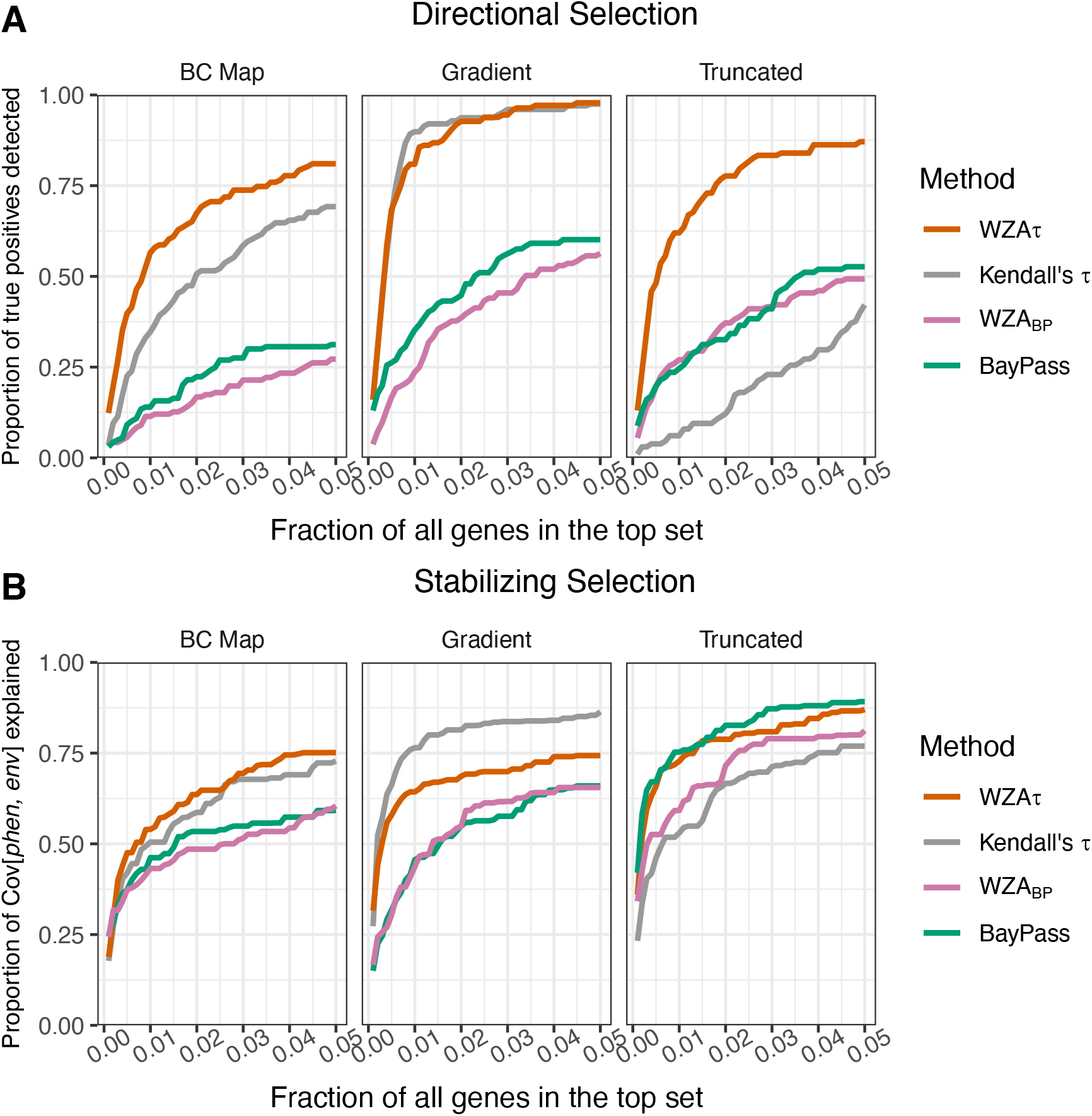
The performance of population structure correction. A) Results for simulations modelling directional selection and b) results for simulations modelling stabilizing selection. Lines represent the mean of 20 simulation replicates where samples of 50 individuals were taken from each of 40 demes. For a description of the x-axis in this plot see the legend to Figure 3.

Notably, even though the Kendall’s τ analysis did not adjust for spatial population structure, the single SNP analyses based on Kendall’s τ in most cases outperformed BayPass (with the exception of stabilizing selection on the *Truncated* map). The discriminatory power of GEAs does not seem to be improved consistently by careful accounting of the underlying pattern of genetic structure.

### The performance of WZA when environmental variables are weakly correlated with selection pressure

In the previous section, we conducted GEA assuming perfect knowledge of the phenotypic optima in each sampled deme. However, environmental variables are often obtained via interpolation and/or may be measured with error, and measured environments may only loosely correlate with the meaningful selective environments. Using the simulations modelling local adaptation on the *BC* map via stabilizing selection, we compared the performance of the WZA against the single-SNP GEA methods when the measured environment is imperfectly correlated with the phenotypic optima.

The WZA outperformed single SNP approaches (Kendall’s *τ* or *BayPass*) when the measured environment was not perfectly correlated with phenotypic optima, especially for weak to moderate correlation between the measured and selective environments (Figure 5). WZA*τ* outperforms the single-SNP approaches when the measurement of the environment is a poor proxy for historical selection.

**Figure 5.**
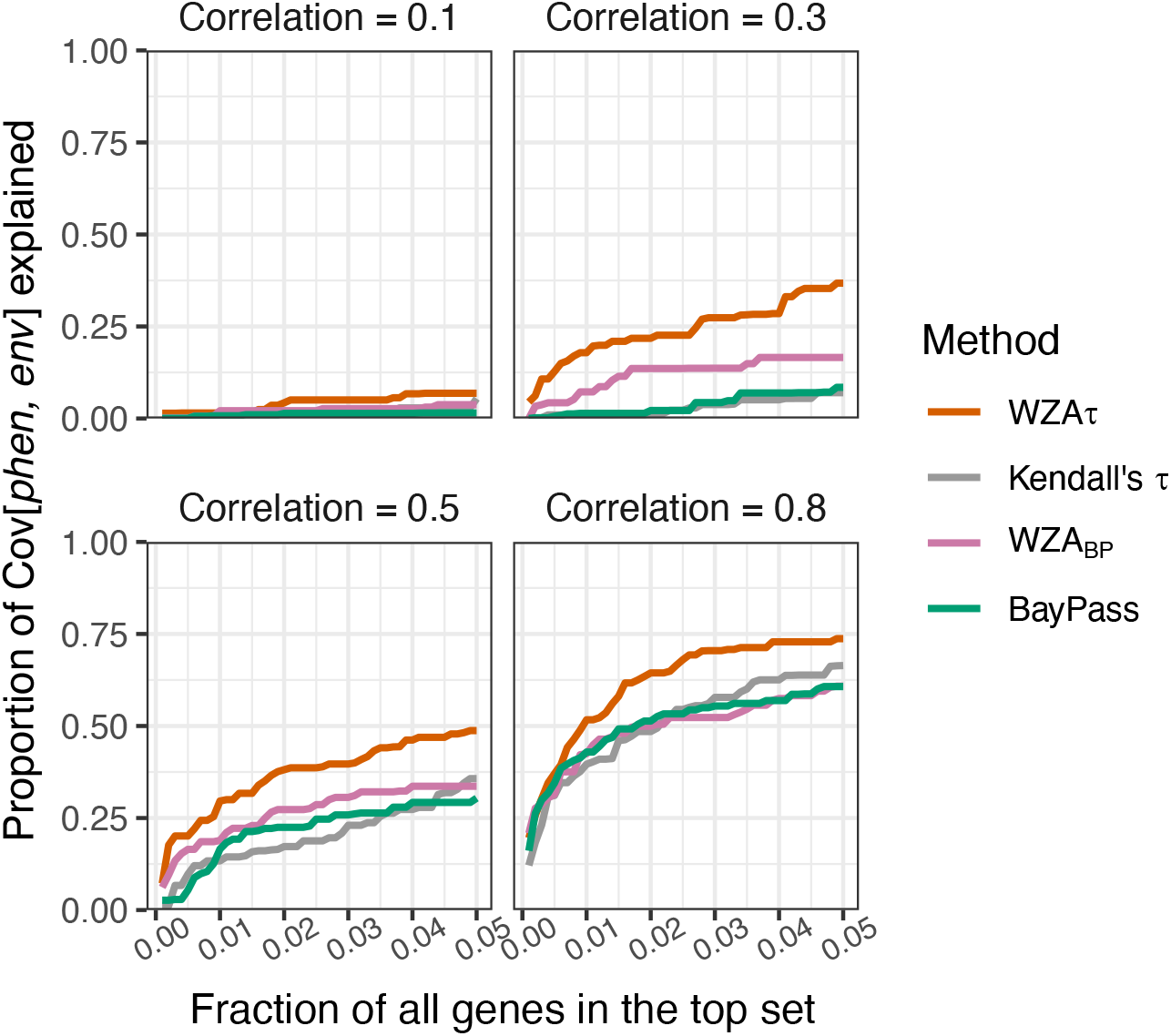
The proportion of true positives recovered when the measured environment is imperfectly correlated with phenotypic optima. The correlation between environment and selection pressure is shown above each panel. Results are from the *BC Map* with stabilizing selection. Lines indicate the means from 20 simulation replicates, and each is based on samples of 50 individuals from each of 40 demes. For a description of the x-axis in this plot see the legend to Figure 3.

### The effect of recombination rate variation on the WZA

Random drift may cause genealogies in some regions of the genome to correlate with environmental variables more than others. Many of the SNPs present in an analysis window that consisted of genealogies that were highly correlated with the environment may be highly significant in a GEA analysis, leading to a large WZA score. This effect would lead to a larger variance in WZA scores for analysis windows that were present in regions of low recombination. To demonstrate this, we down-sampled the tree-sequences we recorded for our simulated populations to model analysis windows present in low recombination regions and performed the WZA on the resulting data. As expected, we found that the variance of the distribution of WZA scores was greater when there was a lower recombination rate (Figure S10). This is a similar effect to that we described in a previous paper focusing on *FST* (Booker et al. 2020).

### Application of the WZA to lodgepole pine data

We re-analyzed a previously published (Yeaman et al. 2016) lodgepole pine (*Pinus contorta*) dataset and compared the WZA to the top-candidate method, which had been developed for the original study. Overall, the WZA and top candidate statistic were broadly correlated and identified many of the same genes as the most strongly associated loci, but also differed in important ways. Across the lodgepole pine genome, there was a mean WZA score of 0.013 with a standard deviation *σ* = 1.67, and a fat right-hand tail (Figure S11). Figure 6A shows the relationship between WZA scores and the –*log*_10_(*p*-value) from the top-candidate method, which were positively correlated (Kendall’s *τ* = 0.245, *p*-value < 10^-16^). When many of the SNPs in a gene had strongly associated statistics, both methods would tend to yield high scores (Figure 6B-C). When there were many SNPs with marginally significant empirical *p*-values (*i.e.* 0.05 < *p* < 0.10) at relatively high frequencies, the WZA method would tend to yield a high score but the top candidate method would not (Figure 6B). By contrast, if the most strongly associated SNPs tended to have low minor allele frequencies, the top candidate method would tend to yield a high score but the WZA would not (Figure 6C). There were several genes that had WZA scores greater than 10 (approximately 6*σ*), but very modest top-candidate scores (Figure 6A). Figure 6B shows that for one such region, there were several SNPs with high mean allele frequency that have small *p*-values. This particular region had a high score from the top-candidate method. Conversely, Figure 6C shows a region that only had a *Z_w_* ≈ 5, but an extreme score from the top-candidate method. In this case, there were numerous SNPs that passed the top-candidate outlier threshold,but they were mostly at low allele frequency. Figures 6C-D show the relationship between allele frequency and the empirical *p*-value for SNPs present in two genes that had extreme scores from both the top-candidate method and the WZA.

**Figure 6.**
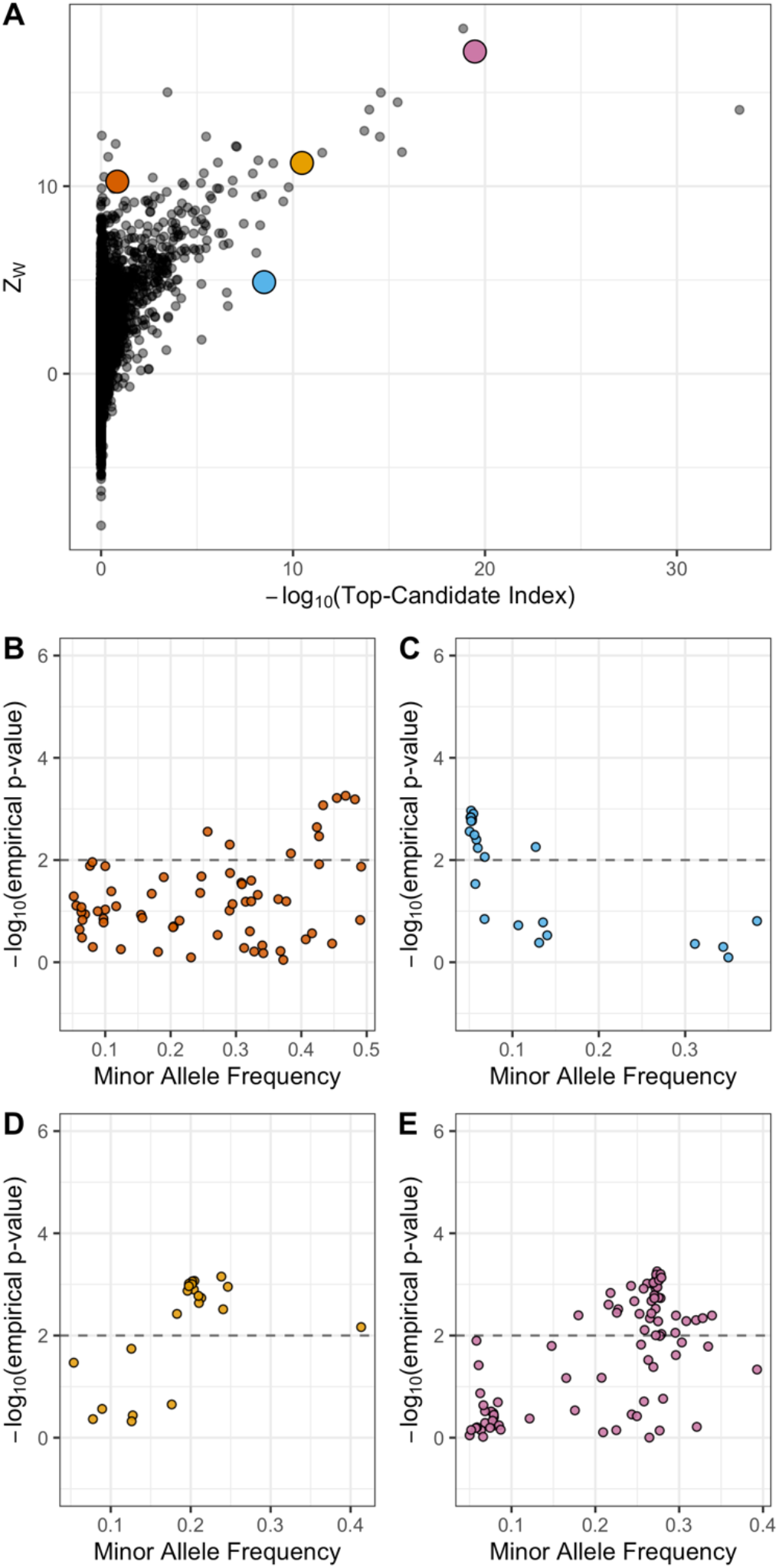
The WZA applied to GEA results on lodgepole pine for degree days below 0 (DD0). A) *Z_w_* scores compared to scores from the top-candidate method for each of the genes analyzed by Yeaman et al. (2016). Panels B-E show the results for –*log*_10_(p-values) for Spearman’s *ρ* applied to individual SNPs against minor allele frequency (MAF) for the colored points in A. The dashed horizontal lines in B-D indicates the significance threshold used for the top-candidate method (i.e. 99*^th^*percentile of GEA –*log*_10_(*p*-values) genome-wide).

## Discussion

In this study, we have shown that combining information across linked sites in GEA analyses is a potentially powerful way to identify genomic loci involved in local adaptation. The method we propose, the WZA, was usually more powerful than looking at individual sites in isolation, particularly when working with small samples or when the environmental variation being analyzed is only weakly correlated with selection (Figures 3 and 5). In a hypothetical world where one had perfect knowledge of allele frequency variation across a species’ range for all sites across the genome, a single marker approach would likely be the best way to perform a GEA analysis, as one would be able to determine the true correlation between genetic and environmental variation for each site in the genome. Indeed, we found that when we had perfect knowledge of allele frequencies in all locations, the SNP-based GEA always outperformed or matched the WZA and top-candidate methods (Figure S12). However, such a situation is unrealistic, and empirical GEA studies will likely always be limited to samples from only some of the populations of interest. Thus, leveraging the correlated information present among closely linked sites in GEA studies may provide a powerful method for identifying the genetic basis of local adaptation.

Theoretical studies of local adaptation suggest that we should expect regions of the genome subject to spatially varying selection pressures to exhibit elevated linkage disequilibrium (LD) relative to the genomic background for a number of reasons. Under local adaptation, alleles are subject to spatial fluctuation in the direction of selection. As a locally adaptive allele spreads in the locations where it is beneficial, it may cause some linked neutral variants to hitchhike along with it (Sakamoto and Innan 2019). LD can be increased further as non-beneficial genetic variants introduced to local populations via gene flow are removed by selection. This process can be thought of as a local barrier to gene flow acting in proportion to the linkage with a selected site (Barton and Bengtsson 1986). Beyond this hitchhiking signature, there is a selective advantage for alleles that are involved in local adaptation to cluster together, particularly in regions of low recombination (Rieseberg 2001; Noor et al. 2001; Kirkpatrick and Barton 2006; Yeaman 2013). For example, in sunflowers and *Littorina* marine snails, there is evidence that regions of suppressed recombination cause alleles involved in local adaptation to be inherited together (Morales et al. 2019; Todesco et al. 2020). The processes we have outlined are not mutually exclusive, but overall, genomic regions containing strongly selected alleles that contribute to local adaptation may have elevated LD and potentially exhibit GEA signals at multiple linked sites. Window-based GEA scans can potentially take advantage of the LD that is induced by local adaptation, aiding in the discovery of locally adaptive genetic variation.

The two window-based GEA methods we compared in this study, the WZA and the top-candidate method of Yeaman et al. (2016), were fairly similar in power in some cases, but the WZA was most often better (Figure 3). Moreover, there are philosophical reasons as to why WZA should be preferred over the top-candidate method. Firstly, the top-candidate method requires the use of an arbitrary significance threshold. This is undesirable, however, because genuine genotype-environment correlations may be very weak and GEA may simply be an underpowered approach to identify alleles that contribute to local adaptation. If there were no detectable signal of local adaptation, ascribing significance to a fraction of the genome may lead to false positives. Secondly, the top-candidate method gives equal weight to all SNPs that have exceeded the significance threshold. For example, with a threshold of *α* = 0.01, genomic regions with only a single outlier are treated in the same way whether that outlier has a *p*-value of 0.009 or 10^-5^. It is desirable to retain information about particularly strong outliers. It should be kept in mind, however, that the WZA (and the top-candidate method for that matter) does not explicitly test for local adaptation and only provides an indication of whether a particular genomic region has a pattern that deviates from the genome-wide average. Indeed, numerous processes other than local adaptation may cause excessive correlation between environmental variables and allele frequencies in particular genomic regions. For example, population expansions can cause allelic surfing, where regions of the genome “surf” to high frequency at leading edges of expanding populations. Allelic surfing can leave heterogeneous patterns of variation across a species range leaving signals across the genome that may resemble local adaptation (Novembre and Di Rienzo 2009; Klopfstein, Currat, and Excoffier 2006).

When performing a genome-scan using a windowed approach a question that inevitably arises is, how to choose the width of analysis windows? If analysis windows were too narrow, there may be little benefit in using a windowed approach over a single-SNP approach. In all the results presented above, 10,000bp analysis windows were used for the WZA. Analysis windows that were narrower than 10,000bp were intermediate in performance between the single-SNP and 10,000bp approaches (Figure S13). Of course, if analysis windows were too wide, the signal of local adaptation may be diluted and the WZA could have little power. It seems like the ideal width for analysis windows would be informed by the pattern of recombination rate variation, LD decay and SNP density across a species genome. In practice, it may be useful to perform the WZA on groups of SNPs, such as genes as in the Yeaman et al. (2016) study. Future study is required to determine the optimal size for analysis windows.

A striking result from our comparison of the various GEA methods we tested in this study was the low power of *BayPass* compared to Kendall’s *τ* (Figure 4). As mentioned in the Introduction, Lotterhos (2019) obtained a similar result in a previous study, though they had used Spearman’s *ρ* rather than Kendall’s *τ*. This presumably occurs because genome-wide population genetic structure is oriented along a similar spatial axis as adaptation, and the correction in *BayPass* therefore causes a reduction in the signal of association at genes involved in adaptation. In such cases, the use of simple rank correlations such as Spearman’s *ρ* or Kendall’s *τ*, which assume that all demes are independent, may often yield a skewed distribution of *p-*values. Such a distribution would lead to a large number of false positives if a standard significance threshold is used (Meirmans 2012). Here, we avoid standard significance testing, and instead make use of an attractive quality of the distribution of *p*-values: SNPs in regions of the genome that contribute to adaptation tend to have extreme p-values, relative to the genome-wide distribution. By converting them to empirical *p*-values, we retain the information contained in the rank-order of *p*-values, but reduce the inflation of their magnitude, which increases the power of the test (Figure S7B). While the empirical *p*-value approach may partially and indirectly correct for false positives due to population structure genome-wide, it loses information contained in the raw *p*-value that represents the deviation of the data from the null model for our summary statistic of interest. It is possible that a GEA approach that produced parametric *p*-values that was adequately controlled for population structure may provide a more powerful input statistic to the WZA, although that was not the case when we tested WZA based on *BayPass* results (Figure 4).

Perhaps more striking is that to identify most or all causal loci, all GEA analyses used here also included a large number of false positives. Previous work has shown that GEA methods are uniformly effective when selection is strong (Lotterhos and Whitlock 2015), so we intentionally simulated weak selection to compare the performance of different methods. Given that we simulated weak selection it is not surprising that GEA methods were underpowered.

Ultimately, performing GEA analyses using analysis windows is an attempt to leverage information from closely linked sites. As mentioned, the WZA could potentially be used with other statistics where LD is expected to result in correlated signals across physically linked nucleotides, for example *p*-values from genome-wide association studies on the basis of phenotypic standing variation, but power in this context would need to be assessed by further testing. With the advent of methods for reconstructing ancestral recombination graphs from population genomic data (Hejase et al. 2020), perhaps a GEA method could be developed that explicitly analyzes inferred genealogies rather than individual markers in a manner similar to regression of phenotypes on genealogies proposed by Ralph et al. (2020). Such a method would require large numbers of individuals with phased genome sequences, which may now be feasible given recent technological advances (Meier et al. 2021).

However, there are scenarios where incorporating information from linked sites in GEA analyses may obscure the signal of local adaptation. For example, the power of the WZA could be reduced if causal alleles contributed to local adaptation along multiple gradients (e.g. to altitudinal gradients in several distinct mountain ranges). If such gradients were semi-independent (i.e. medium/high *FST* among gradients), and then there may be a different combination of neutral variants in high LD with the causal allele in each case. In such a scenario, the species-wide LD in regions flanking the causal locus may be reduced, which would likely also reduce the power of the WZA.

In conclusion, theoretical models of local adaptation suggest that we should expect elevated LD in genomic regions subject to spatially varying selection pressures. For that reason, GEA analyses may gain power by making use of information encoded in patterns of tightly linked genetic variation. The method we propose in this study, the WZA, outperforms single-SNP approaches in a range of settings and so provides researchers with a powerful tool to characterize the genetic basis of local adaptation in population and landscape genomic studies.

## Acknowledgements

Thanks to Pooja Singh for many helpful discussions, to Tongli Wang for help with BC climate data and to Simon Kapitza for help with wrangling raster files. Thanks to Finlay Booker for moral support throughout the course of this project. Thanks to Jared Grummer,Tyler Kent and Isabela Jerônimo Bezerra do Ó for comments on the manuscript. Funding for this work was provided by Genome Canada, Genome Alberta and NSERC Discovery Grants awarded to MCW and SY. SY is supported by an AIHS research. Computational Support was provided by Compute Canada. This study is part of the CoAdapTree project which is funded by Genome Canada (241REF), Genome BC and 16 other sponsors (http://coadaptree.forestry.ubc.ca/sponsors/).

## Appendix

### Parametrizing simulations of local adaptation

Consider a hypothetical species of conifer inhabiting British Columbia, Canada. There may be many hundreds of millions of individuals in this hypothetical species distributed across the landscape. It would be computationally intractable to simulate all individuals forward-in-time incorporating adaptation to environmental variation across the landscape with recombining chromosomes, even with modern population genetic simulators. In our simulations we scaled several population genetic parameters to model a large population when simulating a much smaller one. In the following sections, we outline and justify the approach we used to scale pertinent population genetic parameters.

#### Mutation rate

We set the neutral mutation rate such that there would be an average of around 20 SNPs in each gene after applying a minor allele frequency threshold of >0.05. This number was motivated by the average number of SNPs per gene in the lodgepole pine dataset described by Yeaman et al. (2016). We found that a neutral mutation rate (*μ_neu_*) of 10^-8^ in our simulations achieved an average of 23.3. Note that this *μ_neu_*gave a very low population-mutation rate within demes, 4*N_d_μ_neu_*= 4.0 × 10^-6^.

There are no estimates available of the mutation rate to locally adaptive alleles. We opted to use mutation rates that resulted in multiple locally beneficial alleles establishing in our simulations. For directional selection, we found that a mutation rate of *μ_alpha_* = 3 × 10^-7^ resulted in around 6 locally adaptive genes establishing. For stabilizing selection, a mutation rate of *μ_alpha_* = 1 × 10^-10^, resulted in similar numbers of genes establishing. Note that in our model of directional selection, only a single nucleotide in each of 12 genes could mutate to a locally beneficial allele. In the case of stabilizing selection, all 10,000bp in the simulated gene could give rise to mutations that affected phenotype.

#### Recombination rates

We based our choice of recombination rate on patterns of LD decay reported for conifers. The pattern of LD decay in a panmictic population can be predicted by the population-scaled recombination parameter (*ρ* = 4*N_e_ r*; Charlesworth and Charlesworth 2010), but the pattern of LD decay in structured populations is less well described. In conifers, LD decays very rapidly in conifers and *ρ* ≈ 0.005 has been estimated (Pavy et al. 2012). However, per basepair recombination rates (*r*) in conifers are extremely low, estimated to be on the order of 0.05 cM/Mbp -more than 10× lower than the average for humans (Stapley et al. 2017). This implies a very large effective population size of roughly 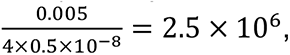 much larger than is feasible to simulate. To acheive a similar number of recombination events through time in our simulated populations, we needed to increase *r* above what has been empirically estimated. We chose a recombination rate that gave us a pattern of LD decay that was similar to what has been observed in conifers. We found that a per base pair recombination *r* = 1 × 10^-7^ (i.e. roughly 200 × greater than in natural populations) gave a pattern of LD in our simulated populations that was similar to what has been reported for conifers.

#### Selection coefficients

It is difficult to choose a realistic set of selection parameters for modelling local adaptation because there are, at present, no estimates of the distribution of fitness effects for mutations that have spatially divergent effects. However, common garden studies of a variety of taxa have estimated fitness differences of up to 35-45% between populations grown in home-like conditions versus away-like conditions (Hereford 2009; Bontrager et al. 2020). Motivated by such studies, we chose to parametrize selection using the fitness difference between home versus away environments.

When modelling directional selection, our simulations contained 12 loci that could mutate to generate a locally beneficial allele. The phenotypic optima that we simulated ranged from −7 to 7 and we modelled selection on a locus as 1 + *s_a_θ* for a homozygote and 1 + ℎ*s_a_θ* for a heterozygote, where *s_a_* is the selection coefficient, *θ* is the phenotypic optimum and ℎ is the dominance coefficient. With a selection coefficient of *s_a_* = 0.003, the maximum relative fitness was (1 + 7 × *s_a_*)^12^= 1.28 for an individual homozygous for all locally beneficial alleles. An individual homozygous for those alleles, but in the oppositely selected environment (i.e. present in the wrong deme) had a fitness of (1 − 7 × *s_a_*)^12^= 0.775. Thus, there would be approximately 40% difference in fitness between well locally adapted individuals at home versus away in the most extreme case. Note, however, that approximately 6 genes established in each simulation replicate, so the realized fitness difference was closer to a 20% difference.

As stated the main text, for stabilizing selection simulations we chose *V*_s_= 192 as this gave a maximum of 50% difference in fitness between individuals grown in home-like conditions versus away-like conditions.

#### Migration rate

We wanted to model populations with *F_ST_* across the metapopulation of approximately 0.05, as has been reported for widely distributed conifer species such as lodgepole pine and interior spruce (Yeaman et al. 2016). For the stepping-stone simulations, we chose a migration rate of 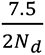 as we found that this gave a mean *F_ST_* of 0.04. For an island model, we used the analytical formulae given in the main text to set *m* to achieve a mean *F_ST_* of 0.03.

---

**Table S1.** Population genetic parameters of a hypothetical organism, and how they are scaled in the simulations. The meta-population inhabits a 14 × 14 2-dimensional stepping stone. Parameters are shown for a population with 12 loci subject to directional selection.

| Parameter | Hypothetical Biological Value | Scaled Parameter | Unscaled (Simulation) |
| --- | --- | --- | --- |
| Global population size ($N_e$) | $10^6$ | - | 19,600 |
| Number of demes ($d$) | 196 | - | 196 |
| Local population size ($N_d$) | 5,100 | - | 100 |
| Recombination rate ($r$) | $2.00 \times 10^{-9}$ | $4N_d r = 0.00004$ | $1 \times 10^{-7}$ |
| Selection coefficient ($s_a$) | 0.0001 | $2N_d s_a = 0.6$ | 0.003 |
| Migration rate ($m$) | $7.35 \times 10^{-4}$ | $2N_d m = 7.5$ | 0.0375 |
| Neutral mutation rate ($\mu_{neu}$) | $2 \times 10^{-10}$ | $4N_e \mu_{neu} = 0.000004$ | $10^{-8}$ |
| Functional mutation rate ($\mu_\alpha$) | $2 \times 10^{-9}$ | $4N_e \mu_\alpha = 0.00004$ | $3 \times 10^{-7}$ |

**Figure S1.**
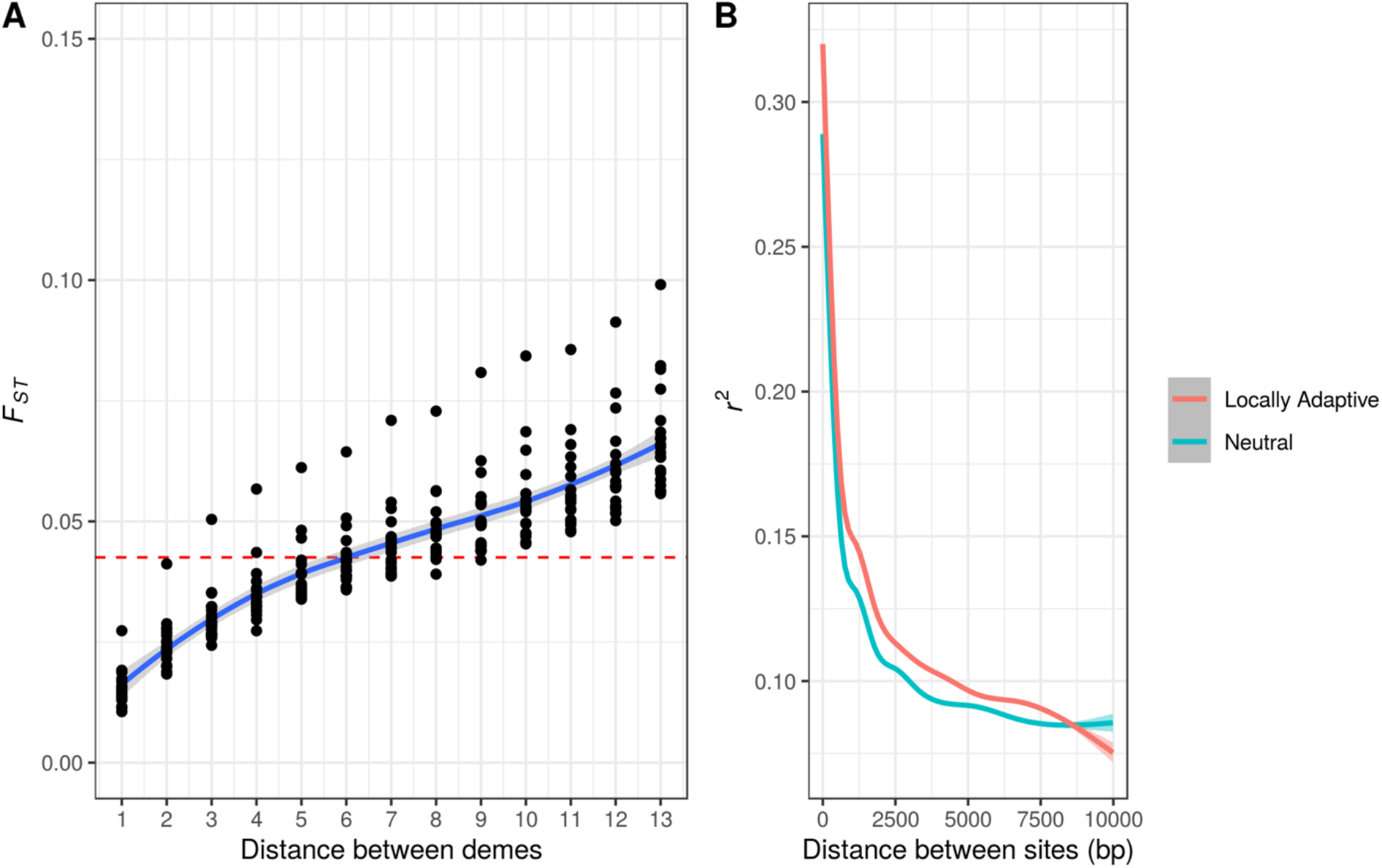
Summary statistics from neutral simulations. A) *F_ST_* between pairs of demes in stepping-stone populations. The average across replicates is 0.042. B) LOESS smoothed LD, as measured by *r*^2^, between pairs of SNPs in genes that are either evolving neutrally are locally adaptation as indicated by the color. Smoothing was performed using the ggplot2 package in R.

**Figure S2.**
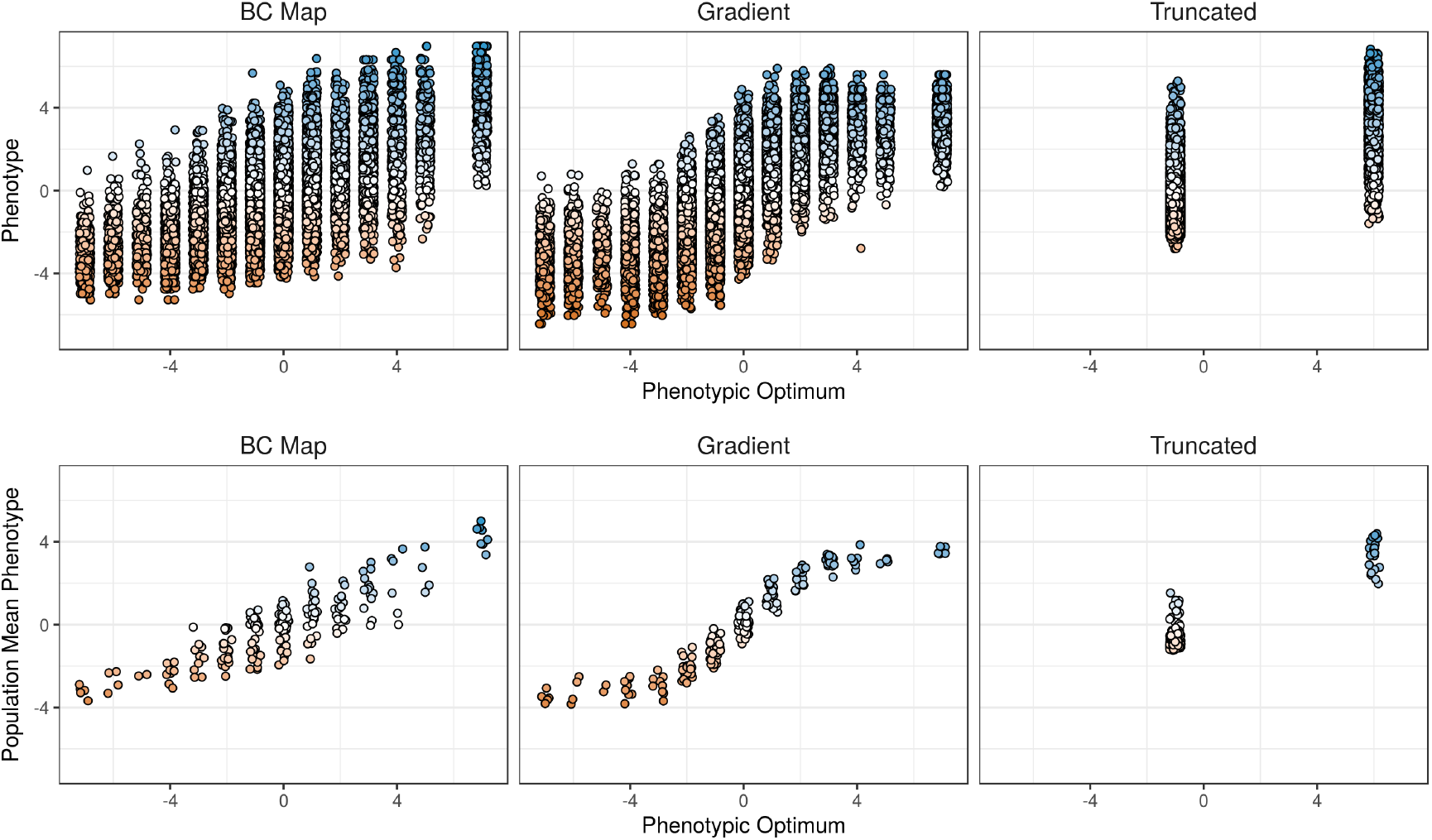
Individual and population mean phenotypes observed in representative simulations for each of the environment maps simulated. A small amount of horizontal jitter was added to points for visualization purposes. Colors represent phenotype values but are for visualization purposes only.

**Figure S3.**
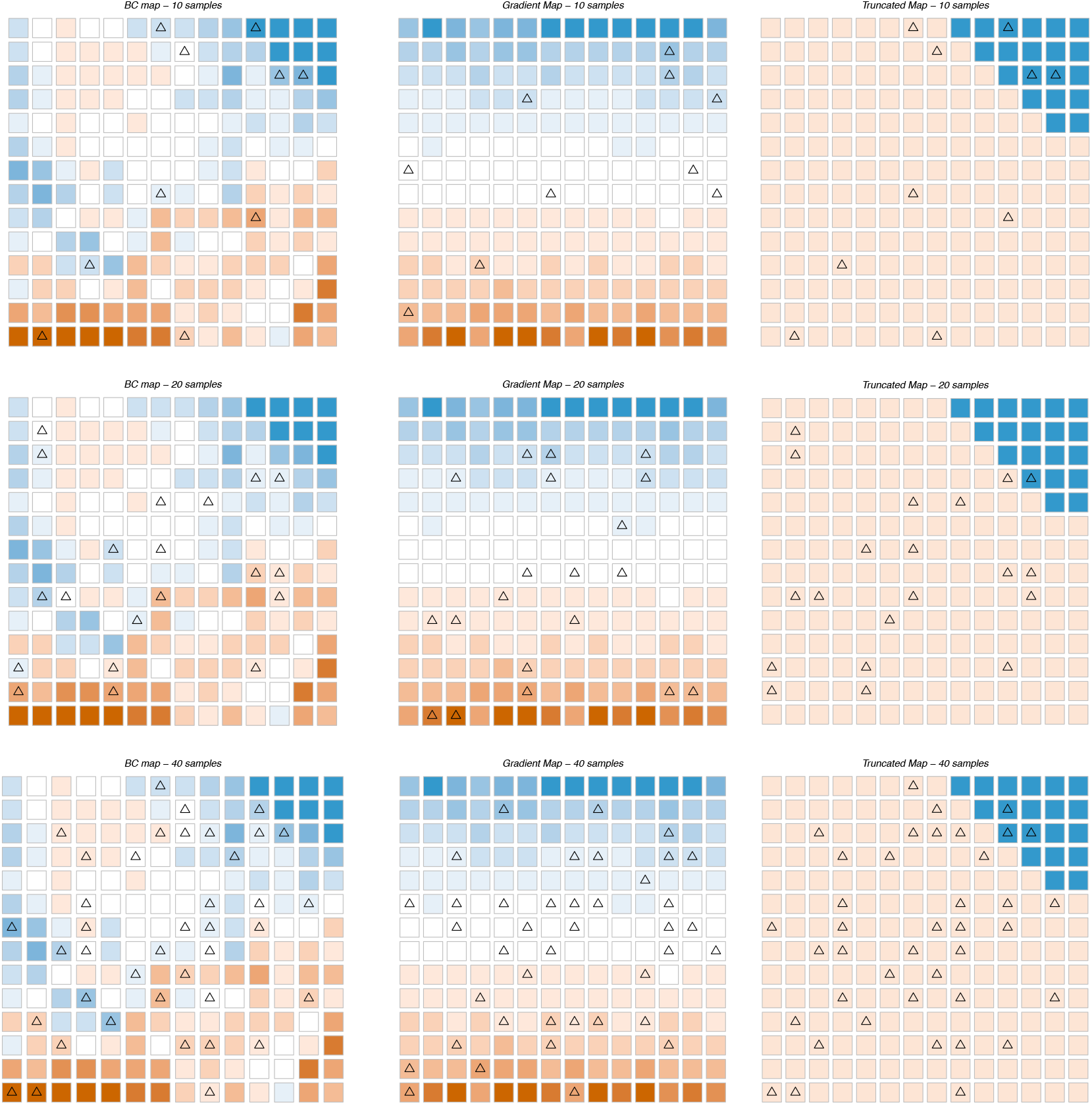
Locations of sampled demes on the maps of environmental variation we assumed in the simulations. Triangles indicate the locations where individuals were sampled in each case. Colors represent the optimal phenotype in each population, using the same color scheme as Figure 1 in the main text.

**Figure S4.**
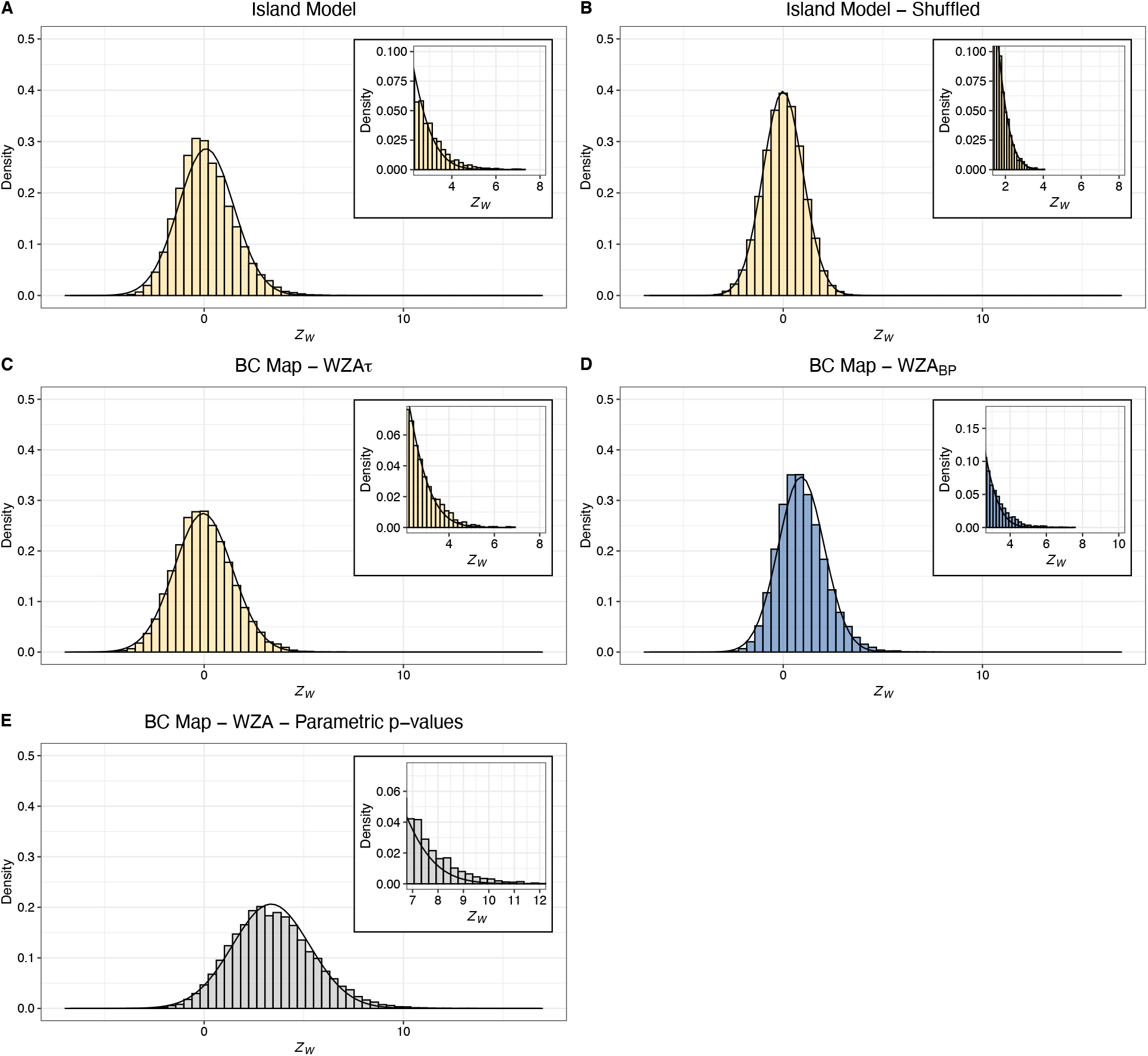
The distribution of WZA scores from neutral simulations with details of the right tail in the insets. Overlaid on each panel is the normal distribution fitted to each dataset. In all cases, results from 20 simulation replicates are plotted together.

**Figure S5.**
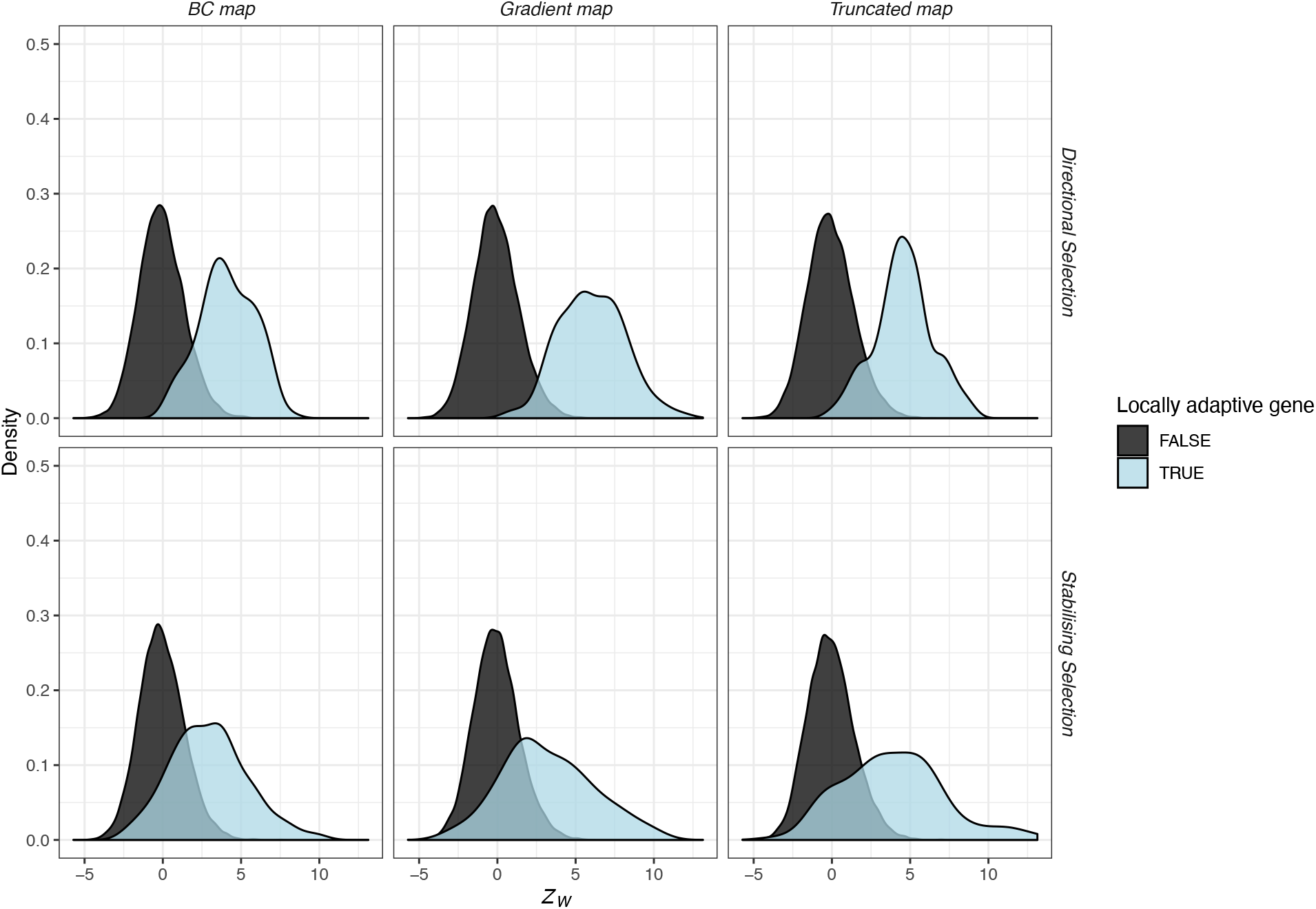
The distribution of WZA scores from simulations of local adaptation. Note, the plot does not indicate the relative frequency of genes that are or are not locally adaptive. Results shown are for samples of 40 demes with 50 individuals sampled in each. In all cases, results from 20 simulation replicates are plotted together. As indicated on the plot, the upper and lower rows contain results for simulations with directional and stabilizing selection, respectively.

**Figure S6.**
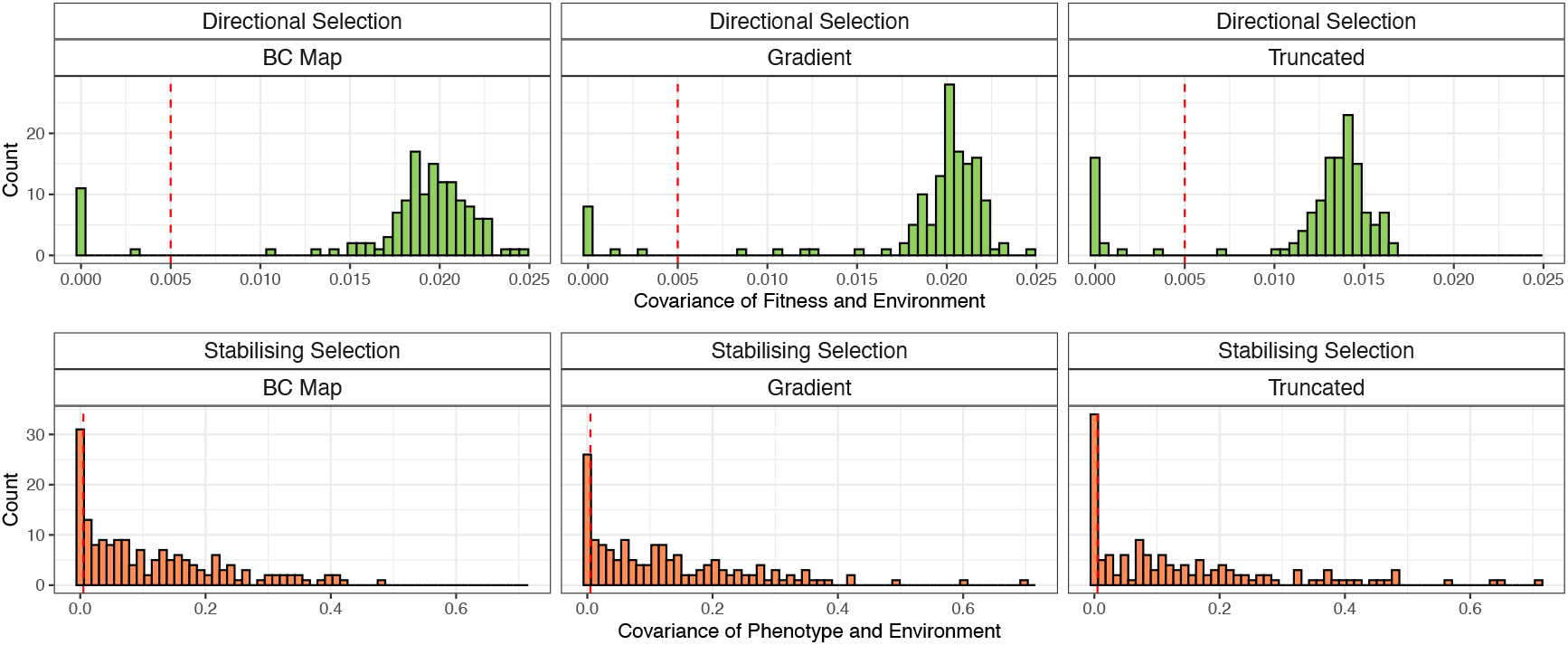
The distribution of effect sizes per gene from simulations of local adaptation. For directional selection, the effect size was we used the covariance between the fitness of a gene and the environment. For stabilizing selection, the effect size was the covariance between phenotypic contribution of a gene and the environment. The vertical line indicates the threshold we applied to the simulated data to classify genes as locally adaptive or not.

**Figure S7.**
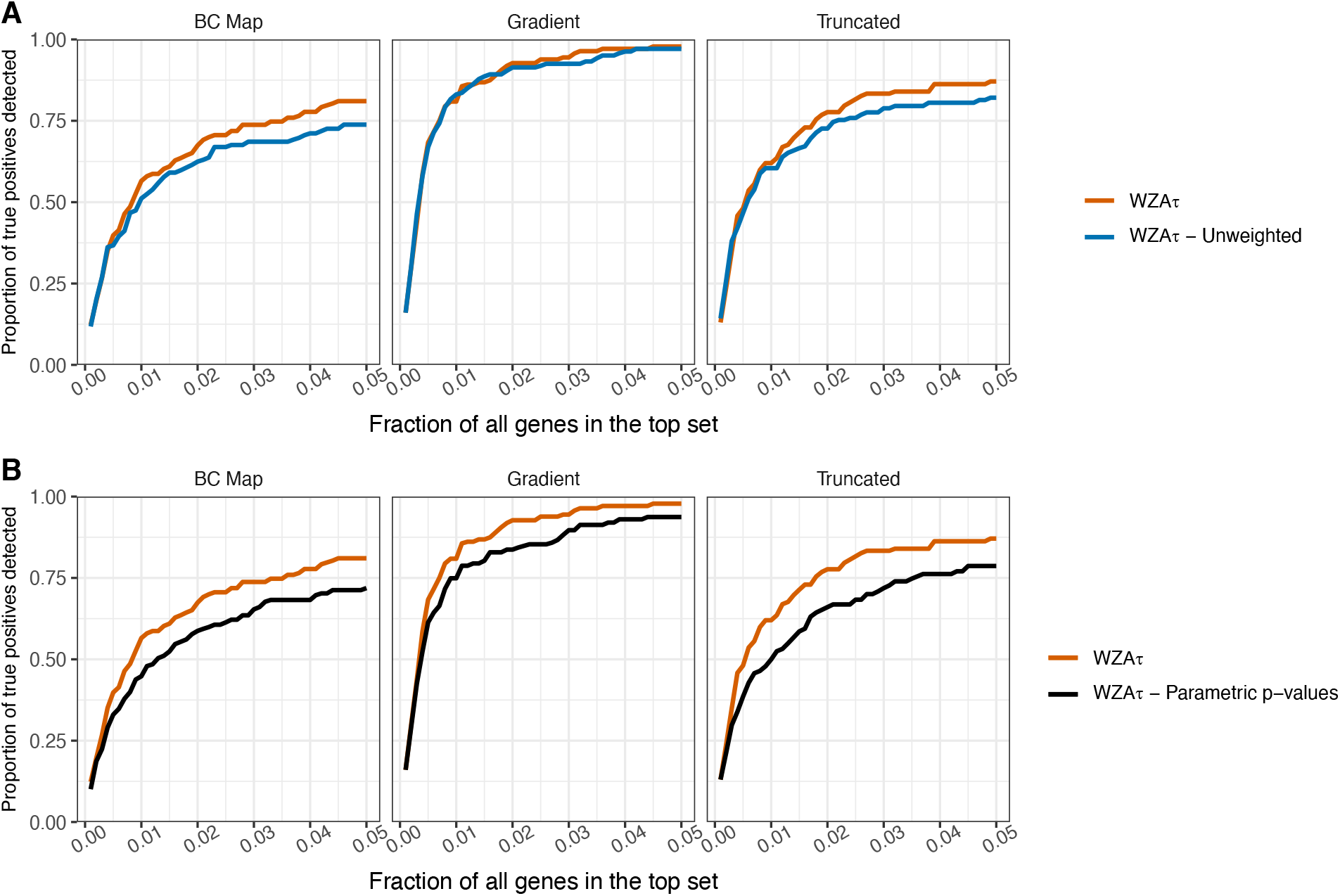
A) Comparison of the WZA performed using empirical *p*-values (WZA*τ*) or using parametric *p-*values from Kendall’s *τ* (WZA*τ* – Parametric *p*-values). B) Comparison of the WZA using *p̅* q̅(as weights in the Equation 1 (WZA*τ*) and an unweighted version of the WZA (WZA*τ* - Unweighted). In each case, the results were obtained using a sample of 50 individuals sampled from each of 40 demes. Lines represent the means of 20 replicates. See the caption of Figure 3 for a description of the x-axis.

**Figure S8.**
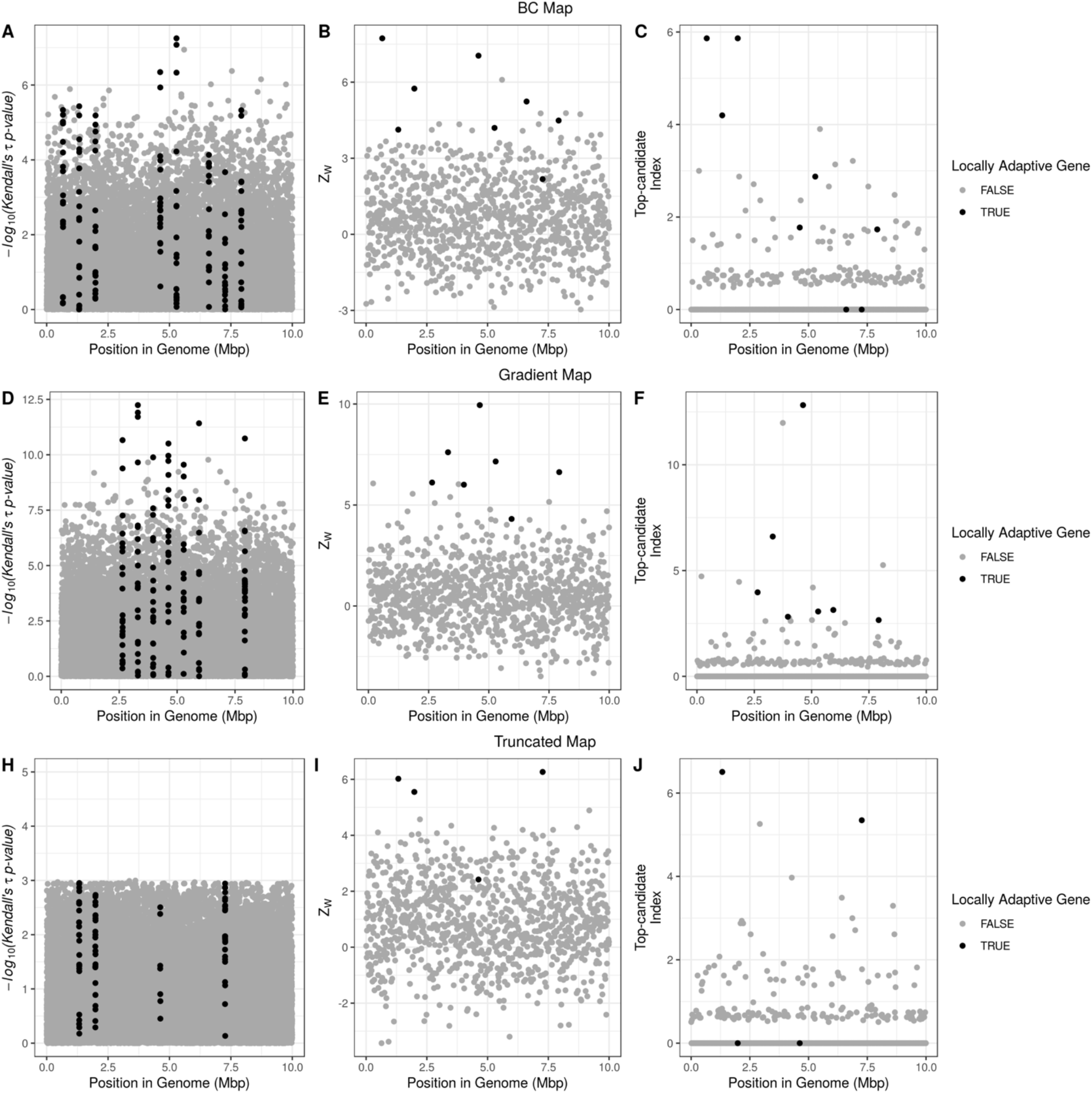
Plots demonstrating the genomic landscape of genotype-environment correlations for a single replicate for each of the three maps of environmental variation we simulated. From top to bottom, the three rows correspond to the *BC Map* (panels A-D), the gradient map (panels E-H) and the truncated map (panels I-L), respectively. The leftmost panel in each row shows the Manhattan plot of –*log*_10_(p-values) from Kendall’s *τ* (panels A, E and I). The central panels in each row show the distribution of *Z_w_* scores from the WZA across the genome (B, F and J) and the distribution of results from the top-candidate method (C, G and K). The rightmost panels show the proportion of locally adapted genes identified using the three different tests for an increasing number of genes in the search effort. Results are shown for directional selection simulations. Note that only SNPs with a minor allele frequency > 0.05 are shown in panels (A, E and I).

**Figure S9.**
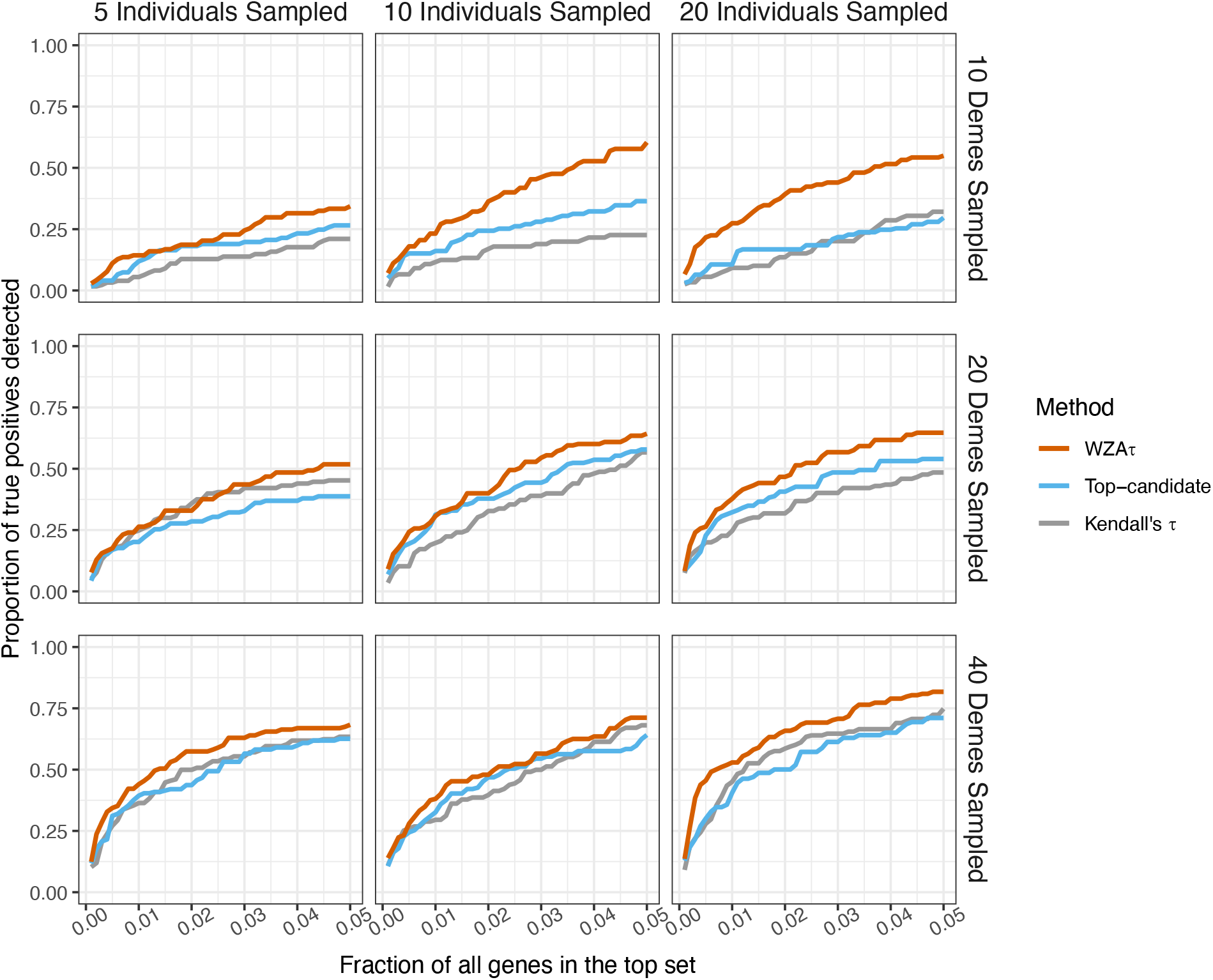
Comparison of the WZA, the top-candidate and the single-SNP approaches with varying numbers of individuals sampled per deme. Simulations shown used the *BC map* and directional selection. Lines represent the mean of 20 simulation replicates. For a description of the axes in this plot see the legend to Figure 3 in the main text.

**Figure S10.**
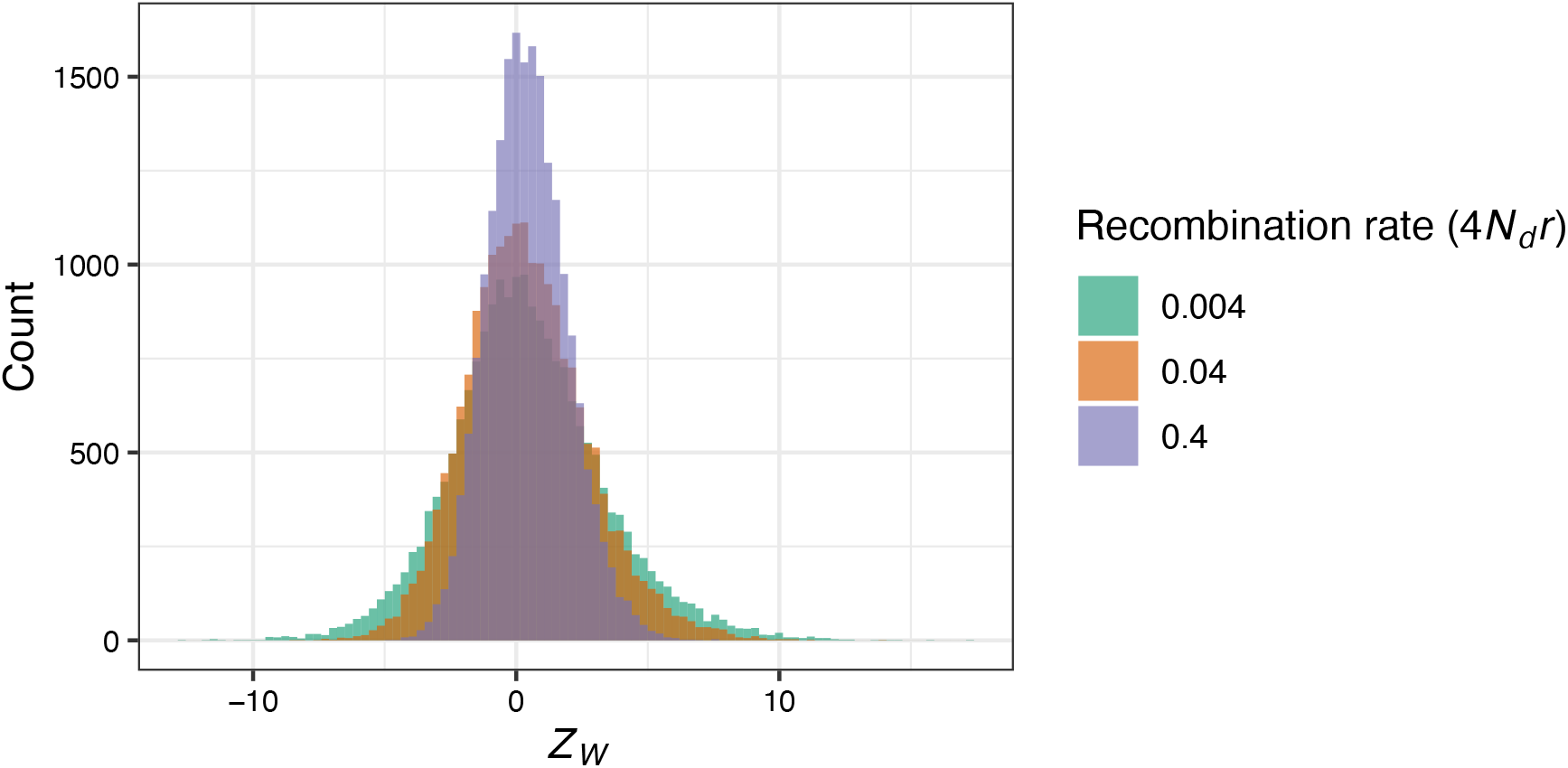
The distribution of *Z_w_* scores under different recombination rates. Results are shown for neutral simulations using the *BC Map.* WZA scores were calculated from a sample of 40 demes where 50 individuals were sampled in each.

**Figure S11.**
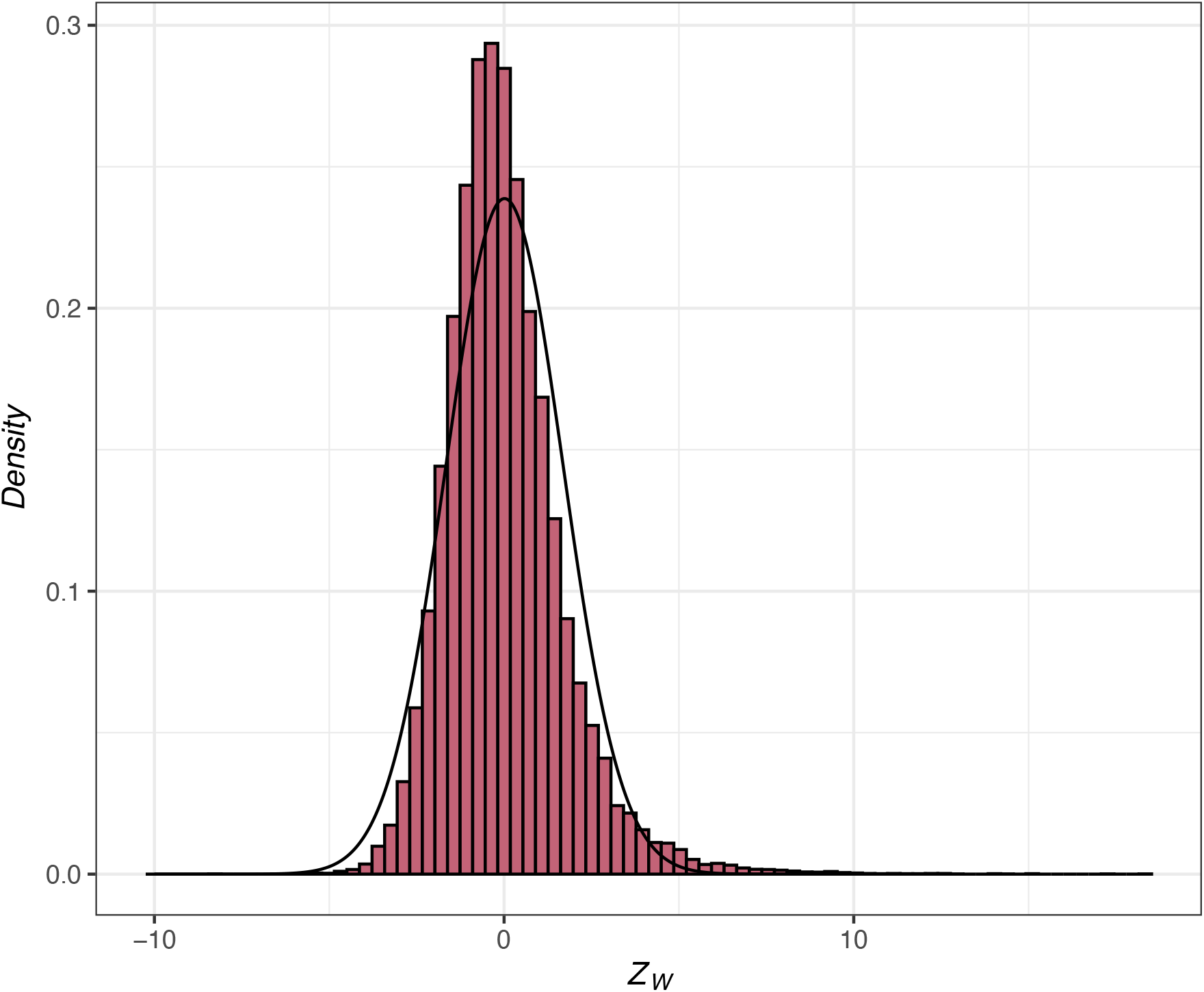
The distribution of *Z_w_* scores for the GEA on (DD0) across the populations of *P. contorta* sampled by Yeaman et al. (2016). The curve shows a normal distribution fitted to the data.

**Figure S12.**
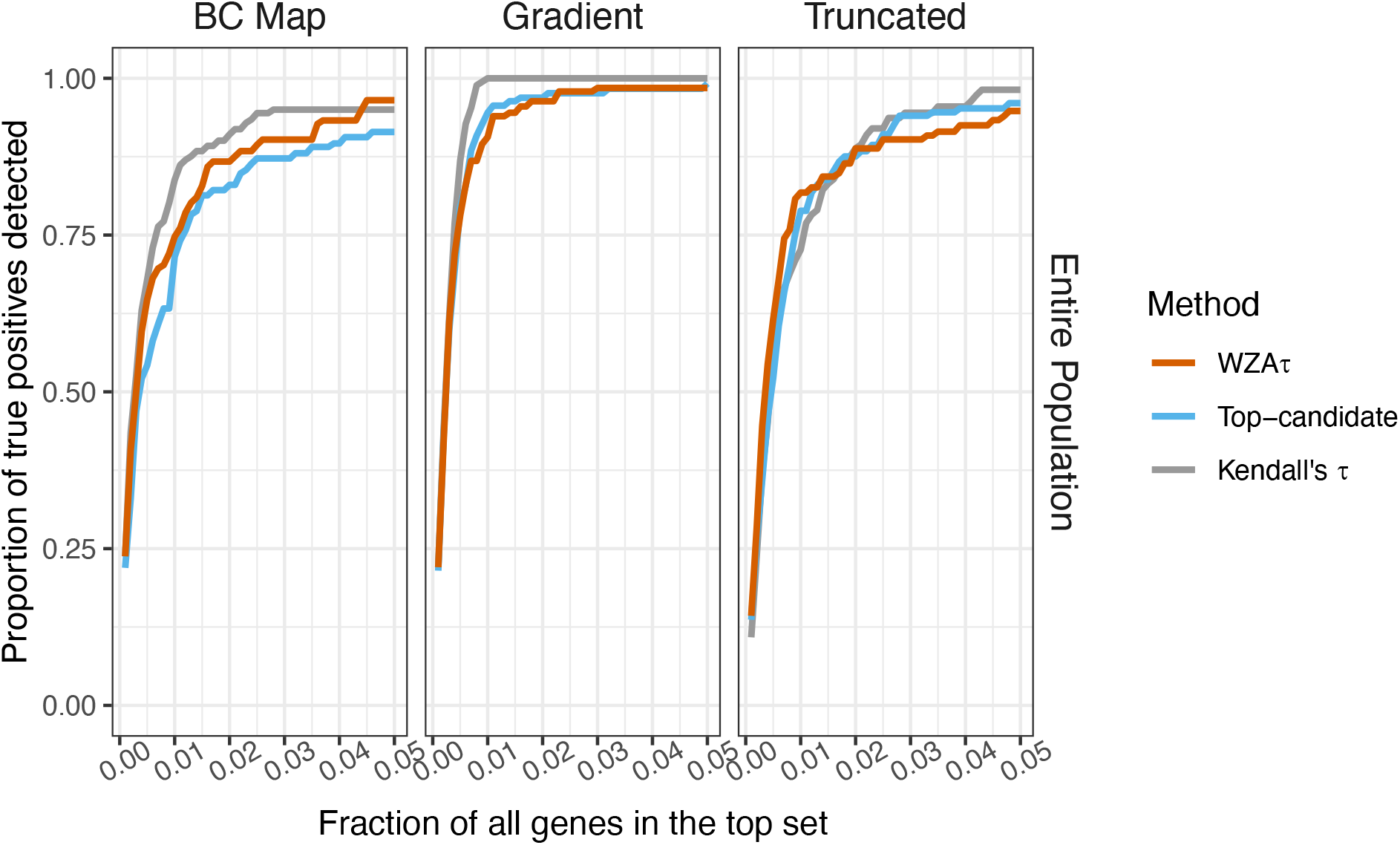
A comparison of three methods to identify the genetic basis of local adaptation when one has complete information on all aspects of the metapopulation, including full sequences for all individuals on all populations. Lines represent the means of 20 replicates. For a description of the axes in this plot see the legend to Figure 3 in the main text

**Figure S13.**
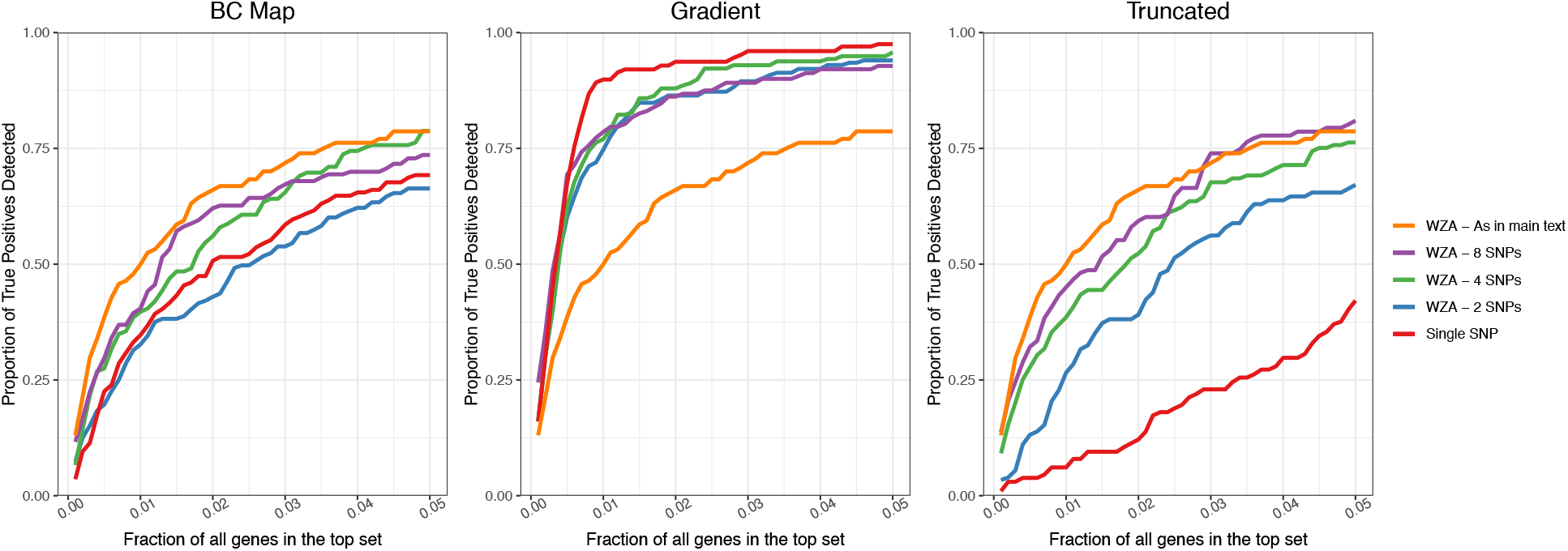
Comparing the performance of the WZA genes identified using the WZA, using analysis windows analyzing a fixed number of SNPs. Lines represent the means of 20 replicates. Analysis was performed on results for a sample of 40 demes with 50 individuals taken in each location. For a description of the axes in this plot see the legend to Figure 3 in the main text

## Notes

### Competing Interest Statement

The authors have declared no competing interest.

